# Oligomeric states of an Influenza-encoded PB1-F2 viroporin

**DOI:** 10.1101/2023.09.09.556987

**Authors:** Sehrish Jamal, Syed Tarique Moin, Shozeb Haider

**Author notes:** Corresponding author: Shozeb Haider.

## Abstract

Influenza Viruses have always been a major health concern due to their highly contagious nature. The PB1-F2 viroporin encoded by the influenza A virus is known to be a pro-apoptotic protein involved in cell death induction of the host immune cells. The structural arrangement and the mode of action of PB1-F2 viroporin have not been fully understood. In this study, we report on the most probable oligomeric structural existences of PB1-F2, investigated by Molecular Dynamics Simulations with improved sampling of conformational states. The simulations provide a channel framework to study the mitochondrial membrane permeation pathway which could initiate the leakage of mitochondrial contents like cytochrome C and induce apoptosis. The structural attributes of the oligomeric states were rigorously evaluated by comparing the experimental reports. Our results reveal a tetrameric form as the preferable state in the lipid environment. This further fulfills the ion transportation criteria by providing a less energetic barrier to ions/water molecules crossing the membrane.

## INTRODUCTION

Influenza Viruses (IV) are highly contagious pathogens that target the human respiratory system. [1]. Influenza virus is an RNA virus that belongs to the Orthomyxoviridae family and is divided into four classes designated as A, B, C, and D depending on their core proteins [2–4]. Influenza A virus (IAV) encodes at least 10 proteins through negatively sensed eight single-stranded viral RNA segments. [5]. In addition, IAV also encodes proteins through frameshifts called accessory or nonstructural proteins [5, 6]. Among those accessory proteins, PB1-F2 is the most discussed protein with pro-apoptotic properties since it kills host immune cells upon viral infection and down-regulates the immune response of the host, particularly through the localization and permeation of the mitochondrial membrane [7] [8]. This protein is expressed only in influenza A-type which is also known to be a potential supporter in boosting epidemics caused by secondary bacterial pneumonia [8] [9–13].

PB1-F2 proteins are encoded by the PB1 gene segment of most of the human and animal IAVs via +1 open reading frame (ORF) [14]. It is a small multifunctional protein that is known for inducing cell death and down-regulating antiviral innate immunity [15–20]. It is also significantly called a promoter of the pro-inflammatory response of cytokines and an active ingredient in the activity enhancement of viral polymerases [5]. Despite the involvement of PB1-F2 proteins in the pathogenicity of influenza, ambiguity persists in its precise role in the influenza life cycle. The structural attributes of PB1-F2 are strain and cell-type-specific but all IAV strains maintain two structural domains which are the N-terminus domain and C-terminus domain [13]. Studies suggest that the protein molecule notably demonstrates a high degree of structural flexibility in the pure aqueous medium, whereas it acquires an extended α-helical structure in membrane mimetic environments [13]. The N-terminus carries two closely-held short helices whereas the C-terminal has a single extended helix, and there is an additional flexible and unstructured hinge region that connects both domains. [13]. PB1-F2 proteins are expressed as 87-90 amino acids, however, truncated N- and C-terminal separates are also common [21–24]. The specialized Mitochondrial Targeting Sequence (MTS) being a part of the C-terminal domain consisting of amino acid residues from Ile55 to Lys85, identified from the synthetic version of protein derived from IAVPR8 targets the mitochondrial inter-membrane space between the inner and outer mitochondrial membrane [13, 25, 26]. MTS then self-assembles to form a positively charged α-helix with clustering of 7 positive regions between amino acid residues from Leu72 to Lys85, and residues Trp58, Trp61, Trp80, and Phe83 holding hydrophobic nature provide structural stability to the protein embedded in the membrane. [10, 18, 25, 26]. Experiments show that PB1-F2 can form channel-like pores in lipid bilayer systems which also supports the concept that these proteins can self-assemble to demonstrate a strong propensity to adopt oligomeric structural arrangements [27]. The protein region largely responsible for oligomerization behavior is located in the C-terminal helix with sequence ^68^ILVFL^72^ and ^54^QIVYW^58^ indicating 93-97%, and 5% contribution, respectively, whereas both, N-terminal and C-terminal domains demonstrate separate oligomerization capabilities [13]. Subsequently, these proteins perturb the mitochondrial membrane potential and facilitate the leakage of mitochondrial contents; for instance, cytochrome C for apoptosis [27, 28] and therefore are characterized as viroporins.

Nevertheless, very limited information is available on PB1-F2 for its viroporin-like behavior, unlike M2 which is considered to be a prototype for viroporins [29–31]. Classically, viroporins are small hydrophobic proteins (approx.100 amino acids) having oligomerization and pore-forming capabilities in membrane environments [29]. An amphipathic α-helix which facilitates the formation of an aqueous channel, and a cluster of positively charged amino acids that anchors the channel to a membrane are the important structural motifs to classify proteins as viroporins [32]. In general, most viruses, for instance, hepatitis C virus, HIV, corona, and picornaviruses, encode at least one of such proteins and avail these specialized molecules for the transportation of ions or other biologically relevant small molecules through the membranes of the host infected cells [33–37]. Viroporins are to a larger extent considered to be expressed by virulent genes as deleting them could disrupt the viral infection cycle [38]. Similarly, the full-length PB1-F2 is also believed to be a virulence factor [9] and has already been known to influence the intensity and the ability to nurture secondary bacterial pneumonia of the pandemic viruses of H1N1-1918, H2N2-1957, and H3N2-1968, along with highly pathogenic avian viruses H5N1 and H7N9, although every influenza strain has a slightly dissimilar PB1-F2 structure [13, 25, 26, 39, 40].

PB1-F2 is also known for its indirect membrane permeabilizing property, for instance by interacting with outer and inner mitochondrial membrane proteins such as Voltage-Dependent Anion Channel 1 (VDAC-1), and Adenine Nucleotide Translocator 3 (ANT3), which constitutes the mitochondrial permeability transition pore complex (mPTP) that could also subsequently induce apoptosis during viral infection [9]. However, the exact mode of action of PB1-F2 at a molecular level has not yet been fully reported except in one study based on molecular dynamics simulations that demonstrated the monomeric structure of PB1-F2 as a non-selective ion channel with truncated N-terminal domain [40].

In this study, we have investigated the oligomerization behavior of PB1-F2 in a realistic membrane environment by employing extensively sampled molecular dynamics simulations. The oligomeric states of PB1-F2 viroporin were modeled in the biomimetic trimeric and tetrameric forms of the recently deposited influenza A virus H1N1 strain. Furthermore, the conformational stability and ion transportation capabilities of both the forms were compared with each other which further aided in understanding the dynamical behavior of the protein and thus establishing its structure-function relationship.

## RESULTS

### Specific sequence motifs may contribute to self-assembly

Over the last 10 years, sequence data isolated for PB1-F2 have shown extreme variation due to rapid and high-level mutations (antigenic drifts) in genetic elements through different events like genetic recombination, horizontal transfer, duplications, and gain/loss of genes [41]. To extract valuable information regarding the possible existence of PB1-F2 in a realistic membrane environment, we aligned the recently reported PB1-F2 protein sequences which were acquired from the major parts of the world, and compared them with the primitively known IAVPR8 strain (A/Puerto Rico/8/1934 H1N1) (Figure 1A).

**Figure 1.**
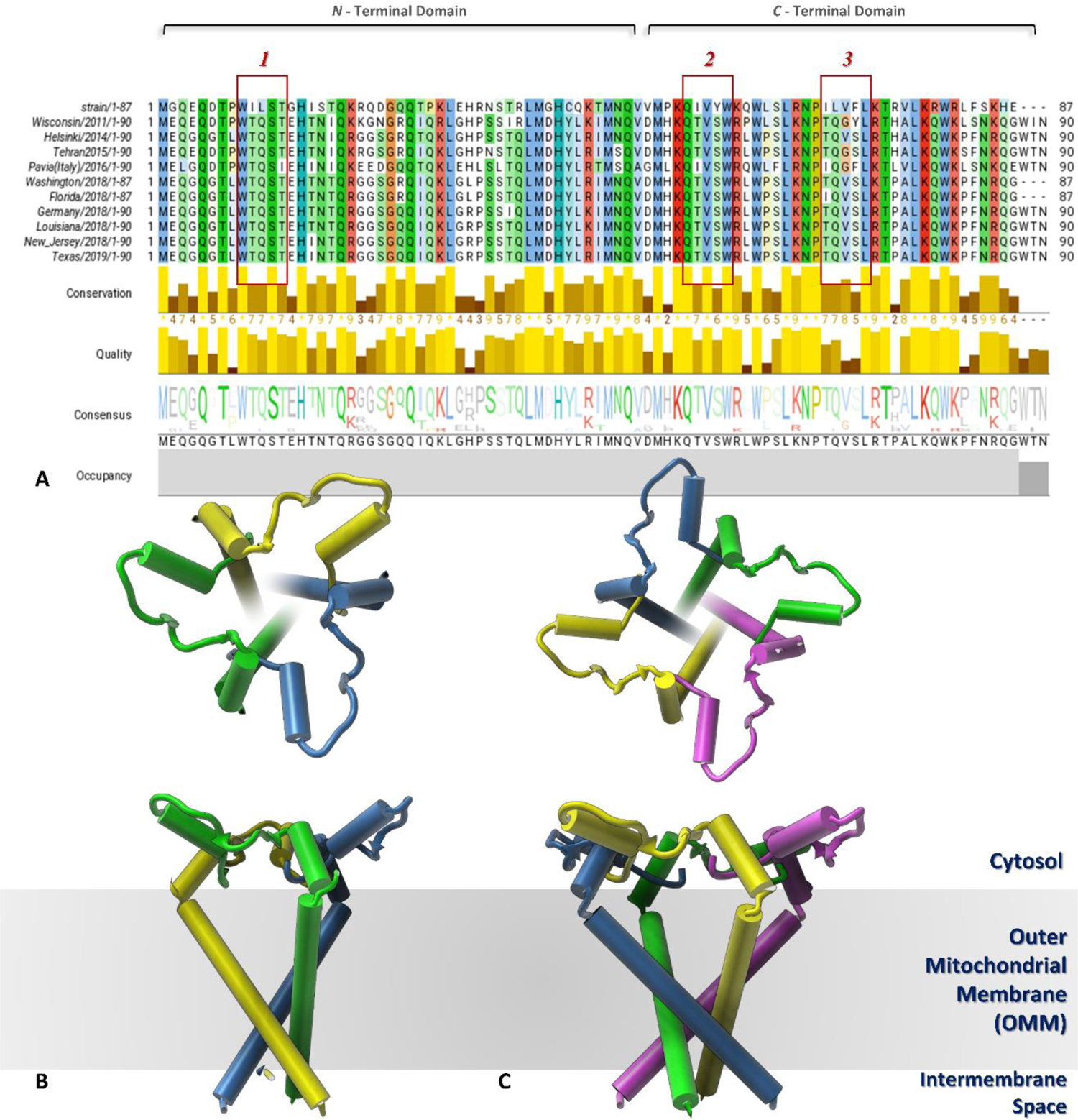
**A**: Sequence alignment of last 10 years of H1N1-PB1F2 isolates with favorable regions for oligomerization highlighted in red boxes; 1 shows the 1%, 2 shows 5%, and 3 shows the 93-97% propensity. **B:** Front and top view of modeled Trimer **C:** Front and top view of modeled Tetramer

The variation in the genetic sequence over time is evident from *Figure 1A* as the neighboring sequences show a greater pairwise identity of ⁓80-90% relative to the sequences placed farther away from each other. However, based on the IAVPR8 strain, motif 1 present in the N-terminal region has an oligomer forming propensity of 1%, motif 2 and motif 3 being in the C-terminal region possess 5% and 93-97% propensity which was also predicted by Bruns et al. in 2007. The greater probability of the 3^rd^ motif for the oligomerization can also be linked to its association with the Mitochondrial Targeting Sequence (MTS). Motif 1 and motif 2 have been observed as relatively conserved, unlike motif 3 which showed variation to specific strains and nevertheless exhibited more capability to alter towards its original pattern. In the current study, we chose the most recent isolate of H1N1-PB1F2 from Texas 2019 and modeled it into trimeric and tetrameric (Figure 1B&C) forms as an extension of Bruns et al. 2007 and Henkel, Mitzner et al. 2010 findings, at the molecular level to investigate the oligomerization properties of this specific protein in detail.

### Assessment of reliable simulation framework

These newly constructed oligomers of PB1-F2 viroporin also supported the findings of Bruns et al. 2007, which described the possession of the C-terminal domain which acquires α-helical conformation with the occupancy of cationic amino acid residues along with the hydrophobic leucine-rich membrane targeting sequence and the solvent-exposed glycine-rich N-terminal domain exhibits random coils or turns like amino acids arrangements thus depicting a true amphipathic nature of the protein. This finding can further be assessed to justify random coil conformations of the viroporin in the presence of an aqueous medium and definite secondary structure (α-helix) in a membrane environment.

Starting from embedding the oligomeric structures of the viroporin in the lipid membrane, the system snapshots were carefully clustered together to extract similar conformations based on the protein RMSDs after carrying out classical molecular dynamics (cMD) simulations. Figure S8 shows the cluster information of each system where the trimer produced a greater number of structural counts along with the first cluster in comparison to the tetramer where simulation time is an exception as more snapshots were considered for the trimer relative to the tetramer. To exclude the possibility of specific structural conformations remaining in local energy minimum upon the simulation treatment, these clusters were essential starting points for further sampling in terms of MIMDS for enhanced conformational space exploration.

### Conformational Stability

The simulation of each oligomeric state of the viroporin demonstrated contrast behavior upon sampling and therefore exhibited diverse conformational changes. By employing the MIMDS approach, each system was sampled through replicates 1 to 10. Both oligomeric systems along with their replicates were passed through the time evolution stability check procedure for the backbone atoms of the domain that is considered to possess important segments in targeting the transmembrane region i.e., membrane targeting sequence (MTS) (Figure S9). The distinct dynamical motions within the same replicates of the trimeric and tetrameric systems authenticate the extensive sampling procedure which was necessary to remove the obstructions that circumvent the structural ensemble to remain in the local energy minimum. The cumulative simulation data was further subjected to compactness analysis for the viroporin’s backbones in terms of the radius of gyration estimations in correlation to root mean square deviations. Figure S10 displays the correlation plots for both oligomers illustrating multiple conformations in the trimeric system whereas reduced conformational space can be envisaged in the tetrameric form thus suggesting a relatively stable system.

The stability of the systems was also analyzed *via* mean square displacement (MSD) of the viroporin along the x-y plane of the lipid bilayers where the extent of translations described the dynamics of replicates in response to the membrane environment (Figure S11). Based on the finding, the conformational drifts in structural ensembles of protein were evaluated to exert distinct effects on lipid molecules due to the presence of a net positive charge towards the bottom of the C-terminal end which could be speculated to drag the protein against the membrane plane and subsequently results in elevated membrane conductances. Besides, the magnitude of the overall displacement was greater for the tetramer than the trimer, thus indicating a significant impact on the geometry of the bilayer composition, and supporting the previous PB1-F2 report regarding mitochondrial membrane short circuits [27].

The oligomerization effects on the surface and curvature of the membrane were also evaluated by considering all the simulation data irrespective of time sequences, therefore the structural ensemble was obtained after an overall sampling time of 2 μs for the trimer and 1.8 μs for the tetramer. The membrane curvature analysis involved the selection of reference surfaces based on the lipid head groups to determine the mean *(H)* and Gaussian *(K)* curvatures of the membranes induced by protein insertion in each snapshot and then averaging all snapshots to produce cumulative curvature plots. As the lipid compositions between each leaflet of the membrane were asymmetric, the calculation was conducted on each leaflet separately. In the case of the trimer, the surface plots of lower and upper membrane leaflets are depicted in Figure 2A. Mean Curvature *(H)* shown in Figure 2B depicted inverted shapes of the surfaces with positive curvature indicating valleys in red color whereas negative curvature corresponded to peaks in blue color. It was evident from the plots that the trimer induced stronger curvature in the upper leaflet in comparison to the lower leaflet. The Gaussian Curvature plots are the measure of the membrane elasticity which indicated that the negative Gaussian curvature represented by pink-colored saddle points and positive Gaussian curvature were shown in green-colored concave regions (Figure 2C). The upper leaflet showed greater flexibility upon insertion of the protein as it had more saddle points and concave regions. This curvature analysis supports the criteria of lipid molecule penetration by the insertion of an ideal viroporin [38] via the attraction of negatively charged lipid phosphate head groups to the cluster of positively charged amino acid residues present in the Membrane targeting sequence (MTS).

**Figure 2.**
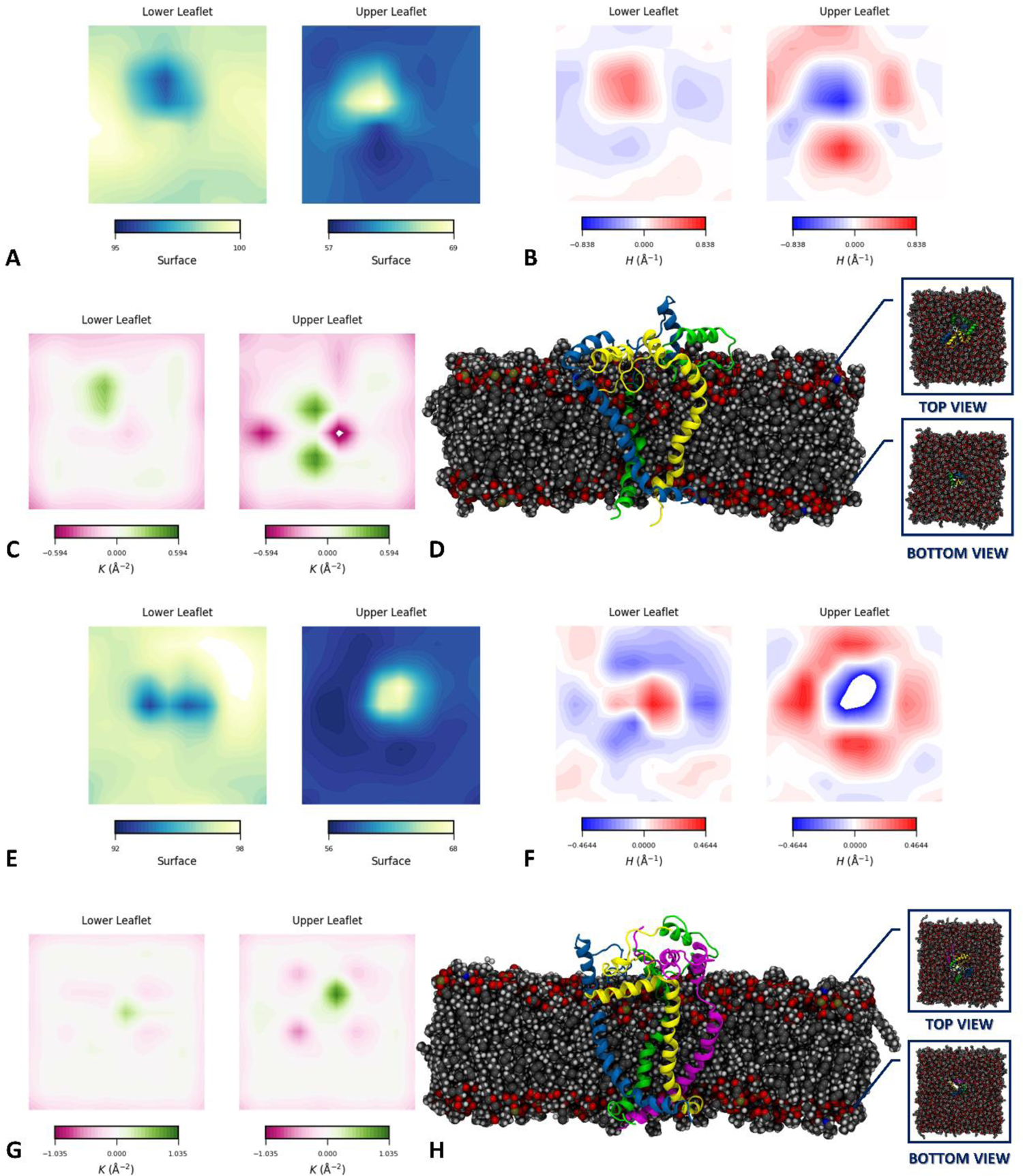
**A, E**: surface plots, **B, F**: Mean Curvature plots, and **C, G**: Gaussian Curvature plots of a lower and upper leaflet of Trimer and Tetramer’s lipid bilayer systems, respectively. **D, H**: Represents the trimer and tetrameric oligomers embedded in a lipid bilayer (gray spheres) with top views (upper leaflet) and bottom views (lower leaflet)

In the case of the tetramer, Mean Curvatures depicted in Figure 2F indicated a more intense effect on the upper leaflet which is most likely due to the presence of an extra homo-oligomeric chain in contrast to trimer. On the other hand, in terms of Gaussian curvatures (Figure 2G), less flexibility can be envisaged relative to the upper leaflet of the trimeric membrane system and that could be an indication of a rigid response upon tetramer insertion into the membrane. Similarly, the lower leaflet also depicted a rigid surface due to the possession of basic amino acid residues at positions K78, K81, and R85 making stabilized contacts with lipid head groups (Table S2)

### Per residual contribution to the overall oligomeric structural states

To assess the residual contribution and behavior of each oligomeric state, replicates were subjected to B-Factor calculations along with Root Mean Square fluctuation (RMSF) pattern estimations for the backbone atoms of both oligomers which revealed homomeric chains to demonstrate distinct dynamical effects on the overall geometry of the viroporin. Considering the case of the trimeric state, the large B-factor and RMSF were observed for the region from S113 to S126 and up to some extent till D130 as this fragment was associated with Chain B N-terminal region (Figure S12). In the case of the tetramer, the terminal M1 amino acid of chain A retained greater mobility in all replicates with random fluctuations showing no regular pattern. However, the tetramer demonstrated major fluctuations in Chain B as observed in the case of the trimer (Figure S13). The high residual fluctuations corresponded to the highly mobile solvent-exposed N-terminal domain in both oligomeric states since the domain contains turns as the major secondary structure element which is assumed to contribute to their instability toward the C-terminal domain linkage that could hold the two domains intact which in result also authenticate the findings related to the presence of random coil or/and unstructured N-terminal segment in an aqueous environment.

To verify residual fluctuations within the homo-oligomeric states of the viroporin, linear correlation metrics were derived from Linear Mutual information (LMI) between the Cα atoms of amino acid residues. LMI was preferred over Dynamical Cross-correlations (DCC) as DCC was insufficient to measure the correlation between two perpendicularly moving atoms which subsequently resulted in a dot product whereas LMI overcame this angular dependency [42]. Figure 3A illustrates the inter- and intra-chain correlations between the amino acid residues of the trimer, and correlations were estimated on a scale from 0 to 1 corresponding to the completely anti-correlated motions and the completely correlated motions, respectively. Based on the matrix plot, the intra-chain residues (depicted with dashed lines) demonstrated distinct behavior within each chain with variable correlative extends between N and C-terminal domains. This correlation behavior indicates the dynamical influence exerted by the neighboring chain’s amino acid residues which can be inferred through highlighted regions from 1 to 4. A strong correlation can therefore be seen between ⁓1-4 amino acid residues of chain C and amino acids residing at the N-and C-terminal interface of chain A depicted as in region 1 whereas strong anti-correlation was evident in region 2 between amino acid residues ⁓20 and 50 of chain C, and ⁓residue 70 of chain A. Amino acid in the region 3 showed anti-correlation between amino acid 19 of chain B and from ⁓20 to 50 of chain C along with a strong correlation between ⁓73 of chain B and ⁓85 of chain C amino acid residues corresponding to the region 4 in the plot. To extract valuable insights from the LMI matrix, centrality analysis was carried out which measures the average links established by each node or more particularly amino acid residues with the neighboring residues since the correlative degree estimations were derived from the LMI matrix of the Protein Contact Network model Figure 3B&C) [43]. Out of the several residues, the estimated highest degree nodes were N84, G87, W88, and T89 of chain A and G87 of chain C (Figure 3C).

**Figure 3.**
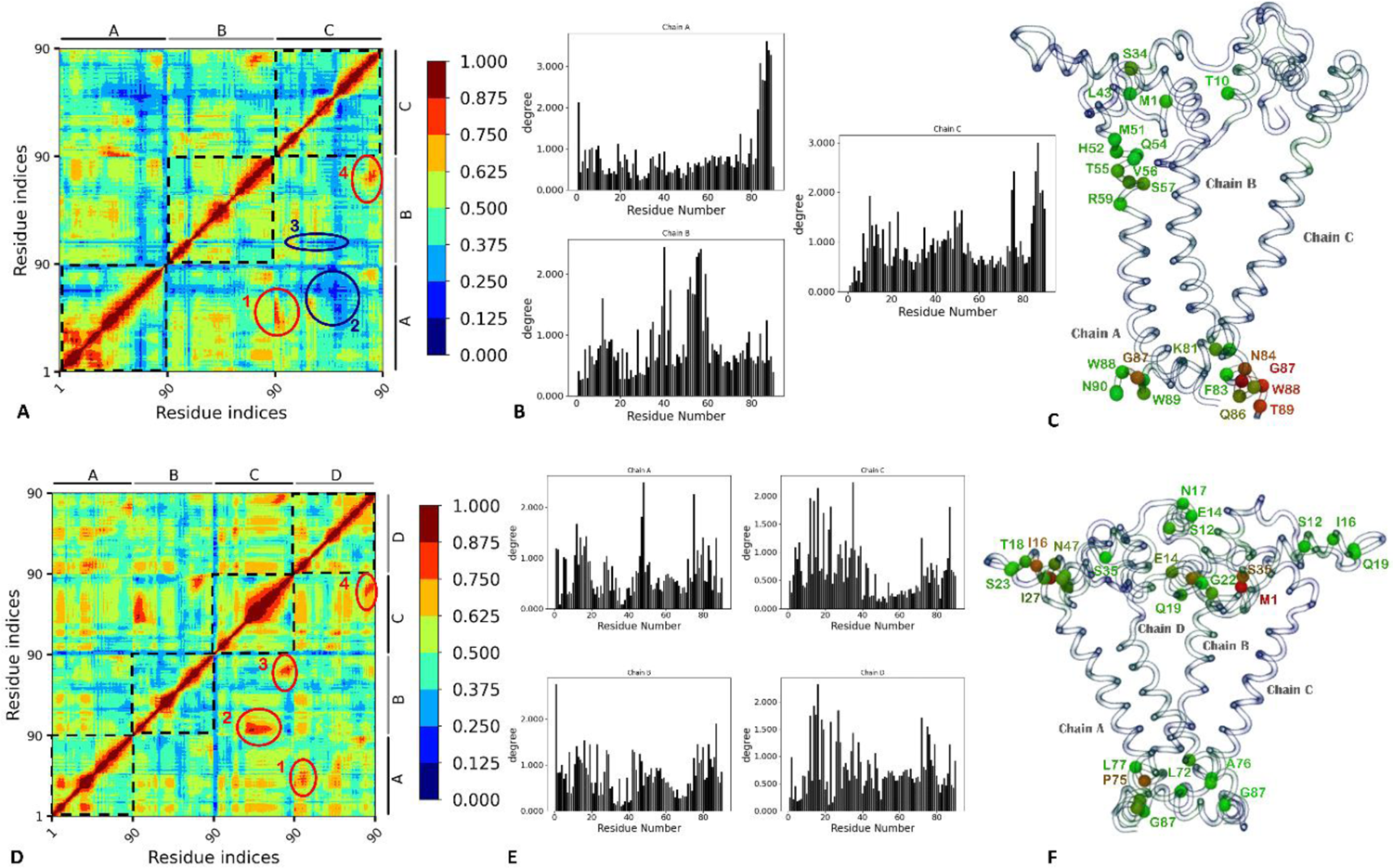
**A, D**: Linear Mutual Information correlation matrix between C-alpha atoms of Trimer and Tetramer, Respectively **B, E**: Correlation Degree plots between the amino acid residues within the chains of Trimer and Tetramer, respectively along with **C, F**: depiction of residual nodes with highest degrees (coloring of spheres is based on BGR scale)

Likewise, the tetramer also showed distinct intra-chain correlation behavior with overall less correlation in chain B as contrary to the trimer case where chain C showed overall less correlation. The correlative motions between the tetramer chains were observed for regions from 1 to 4 illustrated in Figure 3D. Region 1 pointed toward a strong correlation between residues at the interface of the N-and C-terminal of chain A and the first ⁓16 N-terminal residues of chain D. In contrast, residues ⁓37-65 of chain C showed a correlation with the first ⁓19 residues of chain B corresponding to the region 2. Arginine-rich membrane targeting sequence can be seen as having correlated motions between chains C-B which corresponded to region 3, and chains C-D as region 4. The correlative degree plots shown in Figure 3E depicted a similar pattern between N-terminal amino acid residues of chains C and D as possessing greater degree nodes. The prominent residues with the highest degrees were found to be P75, Q48 of chain A, M1 of chain B, I16, S35 of chain C, and I16 of chain D.

The correlated motions between homo-oligomeric chains in the trimer and tetramer were attributed to dynamical responses with each other whereas the tetramer showed more correlative movements in comparison, with more high-degree nodes in the N-terminal region. However, it is also interesting to note that in both oligomeric states charged amino acid residues were not observed in contributing to the highest degree nodes, although polar and hydrophobic residues are mostly involved in stabilizing inter-chain networks which is a characteristic feature of specialized viroporins for oligomeric possessions [32]. Considering the fact that amino acid residues making stable inter-chain contacts showed greater occupancy within the structural ensemble, it could be an effective measure to quantify the elements that retain the viroporin structure in a specific oligomer form. It is therefore, significant to further identify amino acid residues at the interfaces of two or more chains and for this purpose, a cut-off distance of 5.0 Å was considered to probe the inter-chain residual contacts in a simulated structural ensemble. The chain interfaces were examined between chains A, B, and chain C for the trimer, and chains A and B to chains C and D for the tetramer (Figure 4). A detailed interfacial analysis has been tabulated in the supplementary section (*Table S3*).

**Figure 4.**
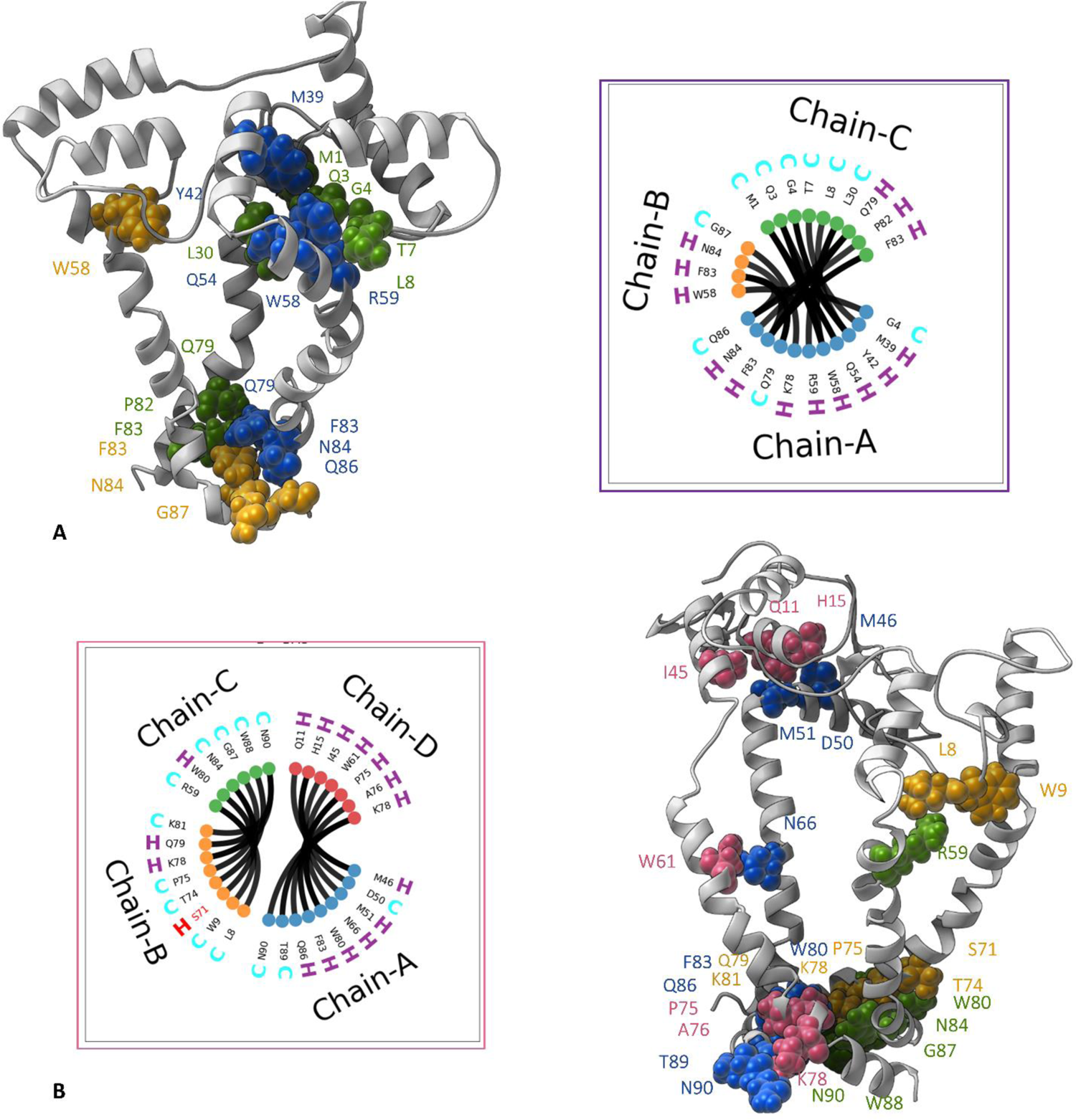
Wheel representations of contacting residues with their secondary structure (H for helix, C for coil) at the chain interfaces with only contacting residues are plotted with cartoon depictions of **A**: trimer and **B**: tetramer form as solid ribbons with interface residues as spheres (color coding based on chain IDs represented in wheel plot)

A brief comparison between interfacial residues of the trimer and the tetramer revealed that the latter contained a large number of interfacial residues in the C-terminal along with an amino acid residue, S71 in chain B that is primitively known from sequence motif 3 (Figure 1A) which was reported to be a favorable region for oligomerization of the protein [13]. However, both oligomers also showed greater contacts between amino acid residues, W58, W80, and F83 which are associated with providing stability toward the membrane. Between the two oligomeric forms, tetramer showed more electrostatically favored interactions which could be assumed to provide structural stability as also evident from contact plots. Within the tetramer form, several inter-residual contacts between chains resulted in linking two monomeric units therefore forming stable dimers authenticating the Bruns et al. findings irrespective of the absence of cysteine amino acid residue which was believed to establish di-sulfide linkages. Furthermore, the solvent-exposed N-terminal domain exhibited random coil conformations and established contacts with amino acid residues of the C-terminal domain at the membrane interface formed between different chains.

Since N-and C-terminal domains have separate oligomerization propensities which can also be deduced from the chain interface and correlation analysis. The residues at the interface linker region of both terminals are expected to play an important role in maintaining the compact helical structure in a full-length state which was reported to be necessary for channel formation [40]. The highly conserved among aligned protein sequences, interfacial hydrophobic residue V49 was further investigated for the contacts between neighboring residues within the cut-off distance of 0.4 nm based on snapshots of all simulation data. The analysis was conducted at every 100^th^ frame after excluding the 4 linearly oriented neighboring amino acids within the same homo-oligomeric chain. Based on the distance distribution plots displayed in Figure S14, the interface residue (V49) in all the chains of both systems shares a neighborhood with the N-terminal residues and has less communication with the C-terminal residues. V49 of Chain A and B in the trimer showed enhanced interaction showing high probability distributions with N-terminal domains. Similarly, V49 of all the chains exhibited increased interaction with the N-N-terminal domain in coil conformation, which was assumed to be favored during the dissociation from the transmembrane domain as reported for other influenza strains existing as truncated PB1-F2 fragments [5]. However, the apparent lower probability distribution in the case of tetramer corroborated the correlation analysis which demonstrated higher degree nodes in the N-terminal region, therefore, establishing strong networks within the protein contact network model.

Additionally, essential dynamics (ED) of the homo-oligomers were evaluated for more meaningful structural manifestations by sorting out the most synchronized protein movements along with the first three principal components. The essential dynamics that contribute towards the conformational space change during the simulation were assigned along the vectors according to the highest variance of the structural distribution [44]. However, the cumulative data irrespective of the chronological time frames were considered, with a stride of 12 and 10 frames for trimer and tetramer, respectively. Subsequently, the high mobility points of both the oligomeric states were classified. The conformational space along the principal components showed maximum transformation with a positional variance of 45.6 % and 44.7 % along with the first principal components of the trimer and the tetramer, respectively. Figure 5A & Figure 5 showed a similar reduction pattern of positional variance therefore rapid reduction of residual flexibility was observed in both systems as the trimer had a slight preference in terms of total flexibility and conformational space. The first eigenvectors of the principal components were further analyzed by plotting the PC1 vs. PC2, PC1 vs. PC3, and PC3 vs. PC2 conformational displacement comparisons where the gradient color from blue to red through white represented the chronological order of conformation scan from the combined structural ensembles of the simulation data. Considering the trimeric state, the projections of MD data along PC1 vs. PC2 (with respective overall variance contribution of 45.63% vs 9.03%) demonstrated two compact clusters centered between planes −80 to −40 and −20 to +40 as depicted in Figure 5A (first upper left panel), and a similar conformational space was observed in PC1 vs PC3 (45.63% vs 6.03%). However, PC3 vs. PC2 (6.03% vs. 9.03%) showed dispersed clusters centered around the −20 to +20 plane. These 2D Projections corresponded to multiple metastable states which were thus corroborated by the Rg-RMSD correlation analysis (Figure S10A).

**Figure 5.**
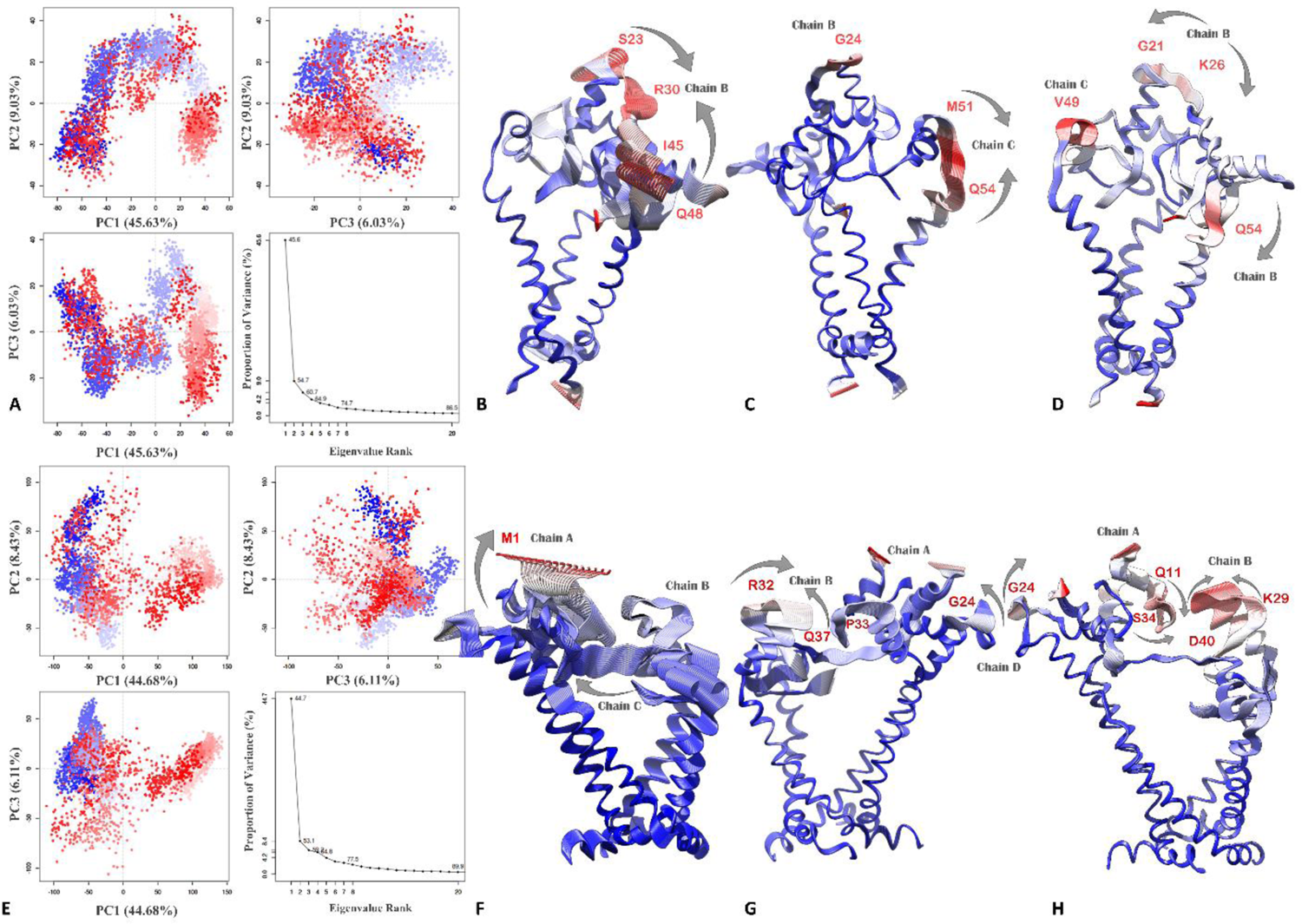
**A**: 2D projections of Molecular Dynamics Trajectories of Tetramer on first three Principal components along with eigenvalue index. **B**: Projected tetrameric conformations along PC1, **C**: PC2, and **D**: PC3, color coding is based on B-factor values (Blue to Red)

Furthermore, each principal component (PC1 to PC3) was examined to identify the major residual fluctuations which were found to encompass the solvent-exposed N-terminal regions in both systems (Figure 5B-D). The highest mobility-inducing residues were from S23 to R30 and I45 to Q48 of chain B which were found along PC1. However, the fluctuations of chain B reduced only to the G24 center with the majority of movements shifted to amino acid residues from M51 to Q54 of chain C, along PC2. PC3 exhibited random motion from G21 to K26 and Q54 of chain B, and intense movement of V49 of chain C. Based on principal components, essential dynamics of trimer were revealed to have occurred in chain B whereas chain A conserved the conformation in the essential sub-space. This confirms the findings of inter-chain residual contact analysis where chains A and C established consistent residual contacts resulting in unconstrained flexibility of chain B up to a certain extent.

Contrary to the trimer, the tetramer exhibited reduced conformational space with a more compact cluster between the −100 to 0 plane in PC1 vs. PC2 (44.62% vs. 8.43 %). In this case, a major grouping was also observed between the −100 to 0 plane, but with greater periodic jumps. PC3 vs. PC2 (6.11% vs. 8.43%) showed a scattered cluster centered between the −50 to +50 plane. Residual fluctuations of amino acid residues along PC1 were randomly distributed with a greater propensity towards the N-terminal domain, and PC2 showed notable motions around R32 to Q37 of chain B, P33 center of chain A, and G24 of chain D. Moreover, intense movements were observed extending from K29 to D40 in chain B and Q11 to S34 in chain A (Figure 5E-H). Overall by comparing the essential dynamics (ED), chain B contributed most of the dynamics in the essential sub-space in both systems despite having a distinct number of homomers. The highly dynamical N-terminal domain in the case of trimer in the essential subspaces also pointed toward the unstable association with the C-terminal domain in the transmembrane region.

### Pore Forming capabilities and Ion permeation electrostatics in the mitochondrial membrane

A very significant aspect of viroporin’s activity is its pore-forming capability in a lipid membrane environment which is thought to facilitate the transportation of ions, water, or any biologically relevant molecules across the membrane [38]. Due to less understanding of the viroporin-like behavior of PB1-F2, as the actual mitochondrial membrane depolarization mechanism of PB1-F2 is still not fully understood, we analyzed the wetting and de-wetting phenomenon of the two oligomeric states to validate the effective transportation of water molecules through the pore. This subsequently helped describe the highly restricted areas or presence of an energetic barrier for water permeation through the channels by profiling the hydrophobic amino acid residues which could play a significant role in the pore formation in the mitochondrial membrane. Therefore, the heuristic prediction analysis was conducted by CHAP (The Channel Annotation Package) [45] on random structural conformations of trimeric and tetrameric states of PB1-F2 to have an insight into water accessibility through the pore, which in turn could help predict the hydrophobic gating mechanism. The water-conductive and non-conductive states of a sampled oligomer structure of the viroporin were predicted by measuring the transmembrane pore dimensions *via* a pore-radius estimation in the presence of a lipid environment derived from the channel’s hydrophobic profiling. The profile defines the maximum and minimum size of the ion permeating or passing through the channel where potential gates were considered as closed if the radius is narrower than the radius of ∼0.15 nm which is a typical value for the radius of a water molecule [46]. The heuristic score is the predictive measure of the channel’s closed gates in a static structure if the sum of shortest distance (Σd) residual points representing as red cubes falling below the dotted line is Σd > 0.55, as shown in Figure S15A&B, the dotted line was separating the wetting (above the line) and dewetting (below the line) of an ion channel. The trimer case represented a barrier for the permeation of water molecules as the computed score was found to be 0.63 with 7 constriction points, whereas, the computed score for the tetramer was 0.56 with 16 constriction points which can be deduced from the plots showing that the tetramer had a slight barrier for water permeation. Moreover, the tetramer possessed a greater pore radius relative to the trimer which showed a large energy barrier for water molecules. This predictive scatch was derived from the profiling of hydrophobic amino acid residues after characterizing transmembrane pore-lining and pore-facing residues (*Table S4*).

Each amino acid residue in a channel structure of the protein was associated with a hydrophobicity value [47] with defined colors for the pore surface visualizations, therefore brown for hydrophobic, blue-green for hydrophilic, and white for neutral amino acid residues, which were plotted (Figure S16) against continuous permeation pathway along the z-axis. The hydrophobicity profiles of trimer and tetramer depicted the pore-lining residues demonstrating hydrophilic properties which could facilitate the transportation of water molecules along the transmembrane channel however the probable localization of hydrophobic lipid side chains in the case of trimer is presumed to be accountable for reduced pore radius [48, 49]. Since the heuristic prediction was based on a single conformation, therefore, to take a deeper look into the permeation pathway, the channel pore radius needed to be frequently calculated from the simulation data (Naz and Moin 2022). For this purpose, the pore-radius was computed for the structural ensemble with the stride of 100 frames by utilizing the HOLE program [50, 51] implemented in MDAnalysis version 2.0.0 [52, 53]. Figure 6A&B showed the variable pore-radius throughout the structural ensemble depicted in multiple color lines with the tetramer having a comparably larger radius than the trimer. The ion specificity of PB1-F2 was also a major concern and was reported to have a non-selective nature, but some reports suggested monovalent ion conduction be favorable for this pro-apoptotic protein [40]. To gain further clarity, we plotted the Poisson–Boltzmann profiles along the channel pores to evaluate the electrostatic environment which is experienced by the particular ions for instance K^+^, Cl^-^, Ca^+2,^ and Na^+^, thus indicating the ion-specific property of the viroporin channel. It appeared as the channels posed a visible barrier for Ca^+2^ ion to a greater extent of 700 kJ/mol for the trimer and 400 kJ/mol for the tetramer, whereas other cations namely, K^+^ and Na^+^ also showed significant barrier in the permeation pathways (Figure 6C&D). In contrast, the ion selectivity was observed for Cl^-^ ions which were more attracted to the side chains of the channel-forming amino acids, such as K65, Q69, L72, R73, A76, Q79, W80, and F83 in both oligomeric states. It was, therefore, presumed that the Cl^-^ ion conduction capability of the tetramer could be enhanced upon changing the pore axis which was derived from the pore-radius calculations as depicted in Figure 6E&F. The radial axis was defined in presence of lipid molecules intrusion thus resulting in constriction points near Q69 and L72 of chain D in tetramer and the trimer with constriction points near L72, R73, Q79, and F83 at C-terminal, and G6, T7, and Q37 at N-terminal with no obstruction for the permeation pathway. This suggested a reliable route for the chloride ion transport across the tetramer which was also justified through heuristic predictions.

**Figure 6.**
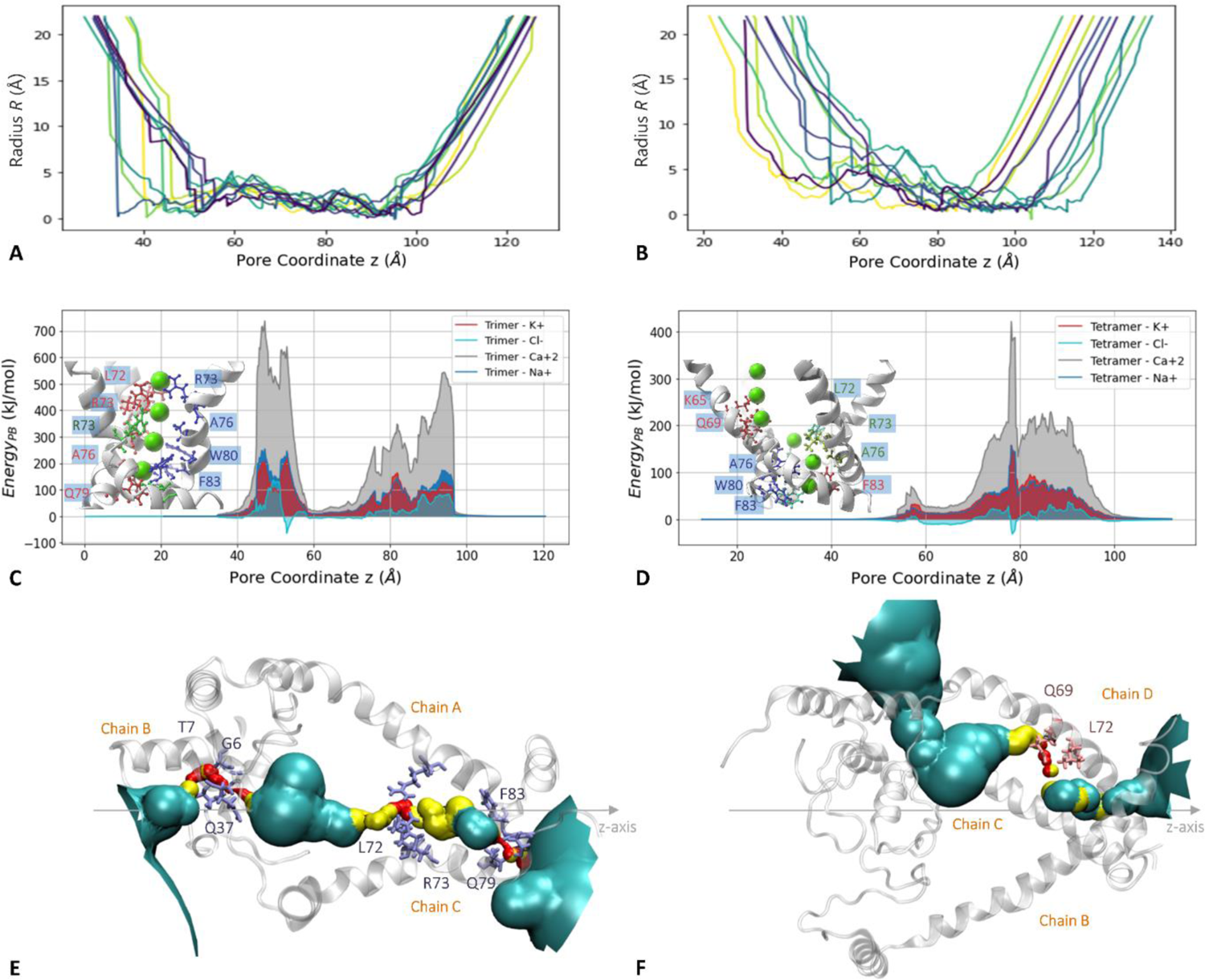
Pore radius of **A**: trimer and **B**: tetramer computed with the stride of 100 snapshots of whole simulation data. Poisson–Boltzmann profiles are along with the **C**: trimer and **D**: tetramer pore with ion-bearing channel depictions. Cartoon representation of constriction points for water/ions transportation along with the **E**: trimer and **F**: tetramer channel z-axis (red surface shows constriction for ions and yellow sphere shows space of maximum to two water molecules

The slight selectivity of the viroporin towards chloride ions is believed to be favored due to the presence of cationic amino acid residues in the membrane targeting sequence, which upon closed examinations of pore radius calculations, were assumed to be not completely accessible due to the intrusion of lipid molecules inside the channel pore. Thus, to characterize the protein-lipid interaction, the simulated structural ensemble was further evaluated. In general, the cationic amino acid residues in specialized viroporin structures are crucial for anchoring the biological membrane via interacting with negatively charged lipid phosphate head groups along with the amphipathic α-helix that facilitates aqueous channel formation [32]. The selected PB1-F2 sequence contains 11 basic/cationic amino acid residues namely lysine and arginine out of which 5 were in the N-terminal segment and 6 were in the C-terminal segment. Since channel-facing basic amino acid residues in the C-terminal domain are of particular interest for anionic conduction, K65 and R73 in each homo-oligomeric chain were monitored for lipid interactions. Figure S17 depicts the percentage occupancy of lipid-amino acid interaction and it is evident that the positive lysine and arginine residues favor the attraction of negative phosphate head groups of lipid making it less available for anion interaction which resulted in a narrower pore radius. The lipid-protein contact analysis represents the consistent interaction of R73 with lipid molecules in the case of trimer whereas K65 of chain A was slightly accessible for non-lipid interactions, on the other hand, in the case of the tetramer, K65 of chain D and R73 of chain A depicted less percentage occupancy which could further support the greater pore radius and lesser heuristic score findings of the tetramer.

## DISCUSSION

The current study provides detailed information on the need for the utmost understanding of PB1-F2, an Influenza A virus-encoded viroporin that has been investigated repeatedly before for its role in the intrinsic apoptotic pathway by disruption of mitochondrial membrane potential, however, the exact membrane permeation insight for probable channel-like and/or lipidic pore formation at the molecular level was still required [27, 40]. PB1-F2 was reported to have a tendency to self-oligomerize and exist in homo-oligomeric subunits through the experimental characterization of the synthetic version of the protein. In this study, the probable oligomeric states (trimer and tetramer) were modeled and embedded in a lipid bilayer by consulting the literature and subjected to an extensive sampling protocol not very often known as MIMDS (Multiple Independent Molecular Dynamics Simulations). The structural attributes of both the oligomeric forms were thoroughly investigated in a membrane environment by utilizing the resultant simulation data for the stability and inter-as well as intra-chain communication analysis followed by ion permeation pathway and lipid accessible site characterizations. At several levels, the obtained molecular detailing was found to augment the experimental findings for instance the oligomerization-favored residues were also seen contributing to stabilizing the specific oligomeric states although, the extent of stabilizing features varied between the two states, with tetramer having more compactness relative to trimer. The effect of trimer and tetramer was also investigated against lipid bilayer in which trimer was found to induce more flexibility to the upper leaflet of lipid whereas tetramer response was relatively rigid despite having an extra homo-oligomeric chain. This behavior presumably fostered by the stabilization factors of W58, W61, W80, and F83 towards the lipid molecules along with the presence of basic amino acid residues (arginine and lysine) as positive counterparts for lipid headgroups. During the simulations, residual contributions within states demonstrated a highly mobile solvent-exposed N-terminal region whereas the C-terminal region possessed correlative motions with greater inter-chain contacts essential for oligomerization behavior. This supported the experimental work by Bruns, Studtrucker et al. 2007, which states the separate oligomeric propensities between the two domains. The tetramer was found to be dominant for inter-chain communication analysis through electrostatic interactions focused on the C-terminal domain. The newly constructed PB1-F2 oligomeric states were also investigated for the evaluation of the hydrophobic gating mechanism through which the amino acid residues in a lipid environment significant for pore-formation were analyzed and predicted to have a greater energy barrier to water for trimer relative to tetramer despite having similar hydrophilic residues along the pore axis, the evident intrusion of hydrophobic lipid side chains was the characteristic behavior that altered the channel profile of the trimeric form. PB1-F2 was previously considered as a non-selective ion channel-forming protein where a slight preference was given to monovalent cations, In this study we evaluated different cations against the permeation pathway out of which all of the analyzed ions showed resistance towards the ion channel except an anion (chloride) which showed appreciable attraction towards the channels apparently due to the positively charged α-helical orientation in the transmembrane region.

## CONCLUSIONS

The study concludes with some major findings which will further aid in unfolding the precise role of the protein, it addresses the poration effect induced by the self-assembly of PB1-F2 and presents the probable structural existence of the protein in a lipid environment. The increased cationic charged density of the viroporin favored its orientation in the lipid membrane in trans due to the apparent accumulation of water molecules and anions which ultimately resulted in the penetration of lipid head groups. This membrane denting behavior could further support the PB1-F2’s lipidic pore formation capabilities more pronouncedly witnessed in the case of the trimer. On the other hand, stabilized strain-specific tetrameric form along with its C- and N-terminal domain portrayed a reliable framework for ion channel activity predominantly due to intense dimeric interactive possessions which could be assumed to adopt improved stability in higher order oligomers for instance octamers. The electrostatic preference for chloride along the channel axis was an essential finding of the study which could be further investigated with longer simulation times with ionic conduction explorations that could also be an effective measure to validate the pore-forming characteristics of the protein either by acting as an ion channel or by promoting the formation of lipid pores, essential for effective ion permeation that subsequently trigger the apoptotic cascade by the release of cytochrome C.

## METHOD

### Sequence Selection and Modeling of Oligomers

The analysis based on the last 10 years of H1N1-IAV strain-specific PB1-F2 sequences was performed by retrieving 11 sequences from the *Influenza Virus Database of NCBI* [54]. This also includes the sequence of the only available 3D structure of the protein (synthetic version) with PDB ID: 2HN8 [13]. After analyzing the pairwise alignments with consensus and conserved regions of protein sequences in chronological order from the date of isolation, a recent sequence with Accession ID QCT08405(A/Texas/9437/2019) was chosen for the modeling of oligomeric forms of the protein.

The full-length experimental 3D structure of PB1-F2 has not yet been reported. The C-terminal domain which is the synthetic version of the protein, displays only 42.11% sequence identity with the C-terminal domain of the selected recent IAV stain. Based on the reported synthetic protein structure, trimeric and tetrameric forms were constructed by utilizing the Galaxy Homomer Suit of the *GALAXY WEB* Server [55]. First, tetrameric homo-oligomer was modeled by overlaying the full-length protein sequence on the template structure with the help of the template-based modeling program *GalaxyTBM* [56]. Next, the modeled tetrameric structure was inspected by consulting the literature which was found to be an appropriate starting framework in terms of the presence of helices and turns. A monomeric subunit from the tetramer model was then taken to yield the trimeric homo-oligomer model by utilizing the *GalaxyOligoTongDock* program which is generally used for *ab-initio* docking treatment that works by implementing M-ZDOCK’s FFT grid-based docking method [57]. The resulting trimeric and tetrameric models were subsequently subjected to *GalaxyRefineComplex* [58] for further structure refinement.

### System Preparation and Classical MD (cMD) Simulations

Both trimeric and tetrameric model structures were embedded in the bilayer system composed of homogenous Palmitoyl-oleoyl phosphatidylcholine (POPC) lipids *via* CHARMM GUI [59] by positioning the Mitochondrial Targeting Sequence (MTS) (amino acid residues ranging from 55 to 85), along the z-axis of transmembrane regions. The trimeric system contained 237 lipid molecules in each leaflet, whereas the tetrameric system contained an unequal number of lipid molecules with 307 POPC molecules in the upper leaflet and 315 molecules in the lower leaflet. Each system was assembled in a rectangular box type with an angle of 90 x 90 x 90° with a system size along the x-y axis of 130 Å for trimer and 150 Å for tetramer. CHARMM36m force field [60] was implemented for the parameterization of proteins and lipids. The lipid-packed trimeric and tetrameric systems were then solvated with the TIP3P water model [61] where 30 Å of water thickness was appended to the ends of the membrane patches, together with the addition of 0.15 M KCl to maintain the neutrality of the systems. Water molecules in the channel pore were retained in both cases where trimer and tetramer held 24 and 109 water molecules, respectively.

Each oligomeric system was then subjected to 5000 energy minimization steps by using the steepest-decent algorithm followed by equilibration in six stages. The first two NVT equilibrations consisted of a total of 250 ps using a time step of 1 fs and positional restraints on protein backbones and side chains with a respective force constant of 4000 kJ mol^-1^ nm^-2^ and 2000 kJ mol^-1^ nm^-2^ applied to the trimer and tetramer system, which were then relaxed to 2000 kJ mol^-1^ nm^-2^ and 1000 kJ mol^-1^ nm^-2^ in the second stage. Similarly, positional restraint was applied to phospholipid phosphorous atoms using a force constant of 1000 kJ mol ^-1^ nm^-2^ which was also relaxed to 400 kJ mol^-1^ nm^-2^, and dihedral restraints from 1000 to 400 kJ mol^-1^ nm^-2^ in the second stage. To maintain the temperature at 303.15 K, the Berendsen weak coupling method [62] was employed. The remaining equilibrations were conducted in an isothermal-isobaric *NPT* ensemble for a total of 375 ps where an initial equilibration aimed for 125 ps equilibration with 1 fs time step and the rest were of 250 ps with the time step of 2 fs. The positional restraints during these stages were relaxed from 1000 to 50 kJ mol^-1^ nm^-2^ for the protein backbones, 500 to 0 kJ mol^-1^ nm^-2^ for the side chains, and 400 to 0 kJ mol^-1^ nm^-2^ applied to the lipid phosphorous atoms, and 200 to 0 kJ mol^-1^ nm^-2^ for the dihedral restraints. The pressure during the NPT equilibration was maintained at 1 bar employing the Berendsen weak coupling method [62]. After equilibration, the systems were subjected to the production stage without restraints, and during the production MD, the Nosé-Hoover thermostat [63] was used to maintain the temperature at 303.15 K with τt (coupling time constant) of 1.0 ps and Parrinello-Rahman barostat [64] for pressure to be maintained at 1 bar with τp (coupling time constant) of 5.0 ps and κp (compressibility) of 4.5 x 10 ^-5^ bar ^-1^. LINCS algorithm [65] was employed to constrain H-bond lengths and the Verlet method was utilized to maintain a buffered neighbor list of all non-bonded interactions with a minimum cutoff of 1.2 nm and PME (Electrostatic Particle Mesh Ewald) [66] to treat long-ranged electrostatics. Short-ranged electrostatics and Van der Waals interactions were also treated with a cutoff of 1.2 nm which smoothly switched to zero from 1.0 to 1.2 nm. Periodic boundary conditions were applied in all directions. The production runs were carried out using the time step of 2 fs for the trimer and tetramer system which were simulated for longer single runs of 1μs and 800 ns, respectively based on the system stabilization time. The all-atom MD simulations were performed using *GROMACS* version 2020.3 [67, 68].

### Clustering and Sampling via Multiple Independent Molecular Dynamics Simulations (MIMDS)

The conformational space of the simulated trajectories was further sampled *via* cluster analysis using the *gmx cluster* tool of GROMACS which implements an rmsd-based clustering approach by utilizing the gromos method [69]. For the clustering of the simulated structures, a cut-off of 0.3 nm was applied on the last 400 ns simulation trajectory of the trimer simulation, whereas a cut-off of 0.4 nm was applied on the last 250 ns of the tetramer simulation trajectory after every 100 snapshots. The specified time spaces were selected based on the stable conformational regions where the RMSD ranges from 0.07 to 0.86 nm for the trimer and 0.09 to 1.31 nm for the tetramer. After clustering, the trimer represented 12 clusters, and the tetramer represented 17 clusters.

The central member from each cluster was extracted whereas the first 10 clusters of each system were considered for further sampling treatment. The Multiple Independent Molecular Dynamics Simulation (MIMDS) approaches [70] was implemented by separately subjecting each cluster representative (10 clusters) to classical molecular dynamics (cMD) production runs of 100 ns with a time step of 2 fs. The simulations of each cluster representative of both systems were conducted using the GPU version of the *pmemd* module implemented in the AMBER20 suite [71] thus the accumulative simulation time for each oligomeric form after clustering was 10 x 100ns = 1μs. Langevin dynamics was employed to control the temperature of the systems with a collision frequency of 1.0 ps ^-1^. Semi-isotropic coordinate scaling for decoupling of z from the x-y plane was applied by utilizing Monte-Carlo barostat where the relaxation time was 1.0 ps and volume change occurred at every 100 steps. SHAKE [72] was applied to constrain all the H-bonds, and long-ranged electrostatics were computed using the particle mesh Ewald (PME) method with force-based switching to treat non-bonded interactions from 1.0 to 1.2 nm. All the cMD simulations were conducted at the isothermal-isobaric *NPT* ensemble.

**Table 1:** Summary of system preparation and MD simulation parameters.

| **System Preparation** | **Parameters** | **Trimer** |  | **Tetramer** |  |
| --- | --- | --- | --- | --- | --- |
|  | *Lipid Composition* | POPC |  | POPC |  |
|  | *No. of Lipids* | Upper leaflet: 237<br>Lower leaflet: 237 |  | Upper leaflet: 307<br>Lower leaflet: 315 |  |
|  | *No. of water molecules* | 45,506 |  | 55,340 |  |
|  | *Total no. of atoms* | 204,716 |  | 255,580 |  |
|  | *System size along the $x$-$y$ axis* | 130 Å |  | 150 Å |  |
| **Molecular Dynamics (MD) Simulations** |  | **Initial Setup** | **Setup after Clustering** | **Initial Setup** | **Setup after Clustering** |
| | *Simulation Time* | $1.0 \mu\text{s}$ | $10 \times 100 \text{ ns} = 1.0 \mu\text{s}$ | $800 \text{ ns}$ | $10 \times 100 \text{ ns} = 1.0 \mu\text{s}$ |

### Trajectory Processing and Data Analysis

All simulation data were analyzed for the acquisition of detailed information on the structural and dynamical properties as well as to have a mechanistic insight into PB1-F2 oligomerization behavior. The data analysis tools used in this study were PyTraj [73], MDAnalysis 2.0.0 [52, 53], Bio3D [44], MDciao 0.0.4 [74], and CHAP [45], whereas the statistical data were plotted by utilizing various modules implemented in Python programming language. Molecular illustrations were generated by Chimera [75] and Visual Molecular Dynamics Software [76].

### Poisson–Boltzmann (PB) Energy Profiles

The energetics of ions in terms of Poisson–Boltzmann (PB) profiles were computed along the channel axis of the trimer and tetramer using the APBS software [77]. The energy profiles were obtained to estimate the electrostatic energy barrier encountered by potassium, chloride, calcium, and sodium ions positioned at specific points along the pore or z-axis derived from HOLE [50, 78] implemented for pore radius calculations. The pore radius estimations were carried out in the presence of a lipid environment to keep the consideration of the lipid sidechain’s interference in the permeation pathway. Partial charges and radii of atoms were calculated using PDB2PQR software [79]. For PB calculations, the systems were positioned in a box having a volume of 97 Å^3^ with the ion specified at a certain distance. The number of grid points was 100 x 100 x 105 Å whereas the fine grid points to focus the placed ions were 10 Å^3^. The dielectric constant was 2.0 and 78.5 for protein and solvent, respectively, and a probe radius of 1.4 Å was considered for the solvent, whereas the concentration of mobile NaCl ions was 0.15 M. The PB calculations for the ion were carried out at every instance of its occurrence in the pore, whereas the charges and born radii of K^+^, Cl^-^, Ca^+2^, and Na^+^ were specified as +1*e* and 2.172 Å, −1*e* and 1.937 Å, +2*e* and 1.862 Å, and +1*e* and 1.680 Å, respectively [80]. The PB calculations were carried out simultaneously by placing each ion along the ‘z’ coordinate according to the following equation.

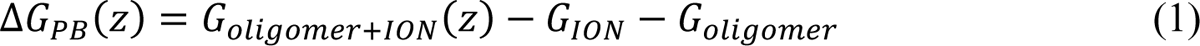

## SUPPLEMENTARY INFORMATION

### Structure Quality Check analysis

Quality assessment of newly constructed oligomers was an essential requirement to gain insight into structural properties, therefore we evaluated the quality of the modeled proteins by plotting the Ramachandran Plots of a monomeric unit of the protein (Figure S7).

**Figure S7.**
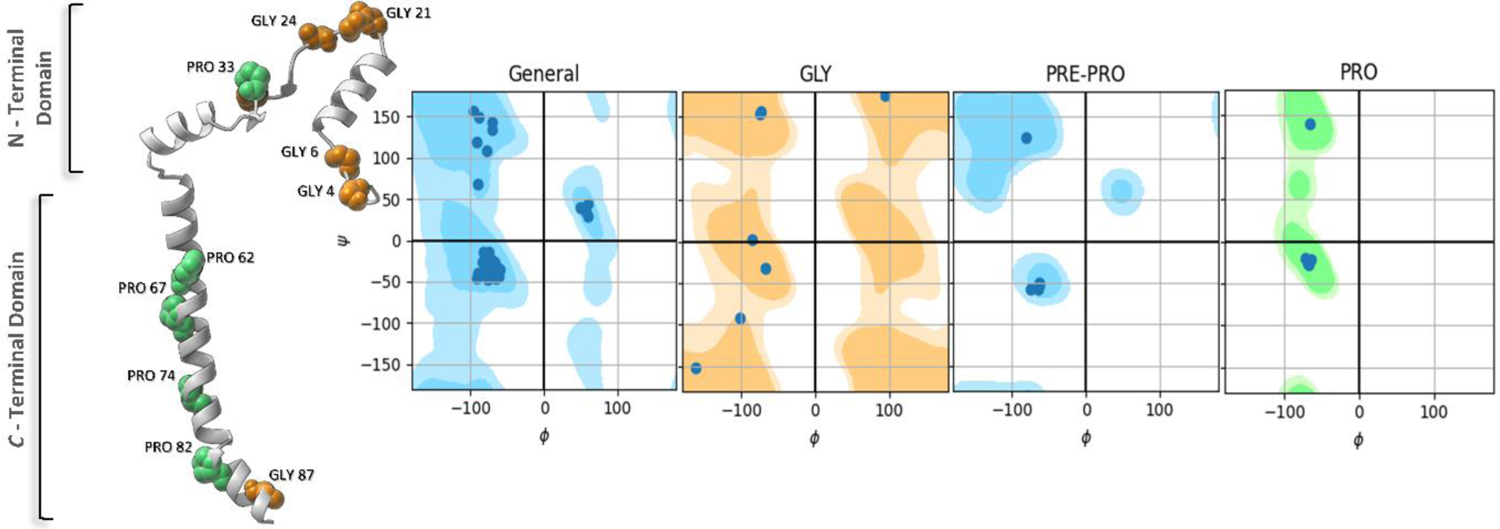
Ramachandran Plots of Monomeric unit with cartoon representation where Glycine residues are highlighted in orange and Proline residues in green color spheres.

Figure S7 shows phi (φ) and psi (ψ) dihedral angles of amino acids like glycine, pre-proline, and proline which were reported to be the characteristic predictors for the secondary structure validation of any protein. The dark color contours in the plots depict the favorable regions with no steric clashes whereas the light-colored regions show the allowed regions. It is evident from the plots that the newly constructed protein represents a good quality secondary structure as no residue appeared in the disallowed region and has a greater propensity towards right-handed α-helix conformation in contrast to few residues in the β-stranded and left-handed α-helix regions. It is pertinent to mention here that glycine and proline are often separately considered in designating a phi (φ) and psi (ψ) dihedral angle due to their helix-breaking nature. Glycine possesses greater flexibility due to the lack of a sidechain that could form bonds with other amino acids and therefore it is considered highly uncertain in designating dihedral angles. On the other hand, proline lacks the hydrogen on its side chain nitrogen atom, and consequently, it is unlikely to form hydrogen bond interaction with other amino acids which are necessary for adopting helical conformations thus creating a kink if located in between a long helical region. Further, it has also a restricted rotation around the N-Cα bond which produces a steric clash to the side chain in the next turn of the helix resulting in 30° bending in the helix axis. Due to such stereochemical behavior, glycine and proline are often involved in the formation of turns which can be observed in the cartoon representation in Figure S7.

### Clustered Conformations

**Figure S8.**
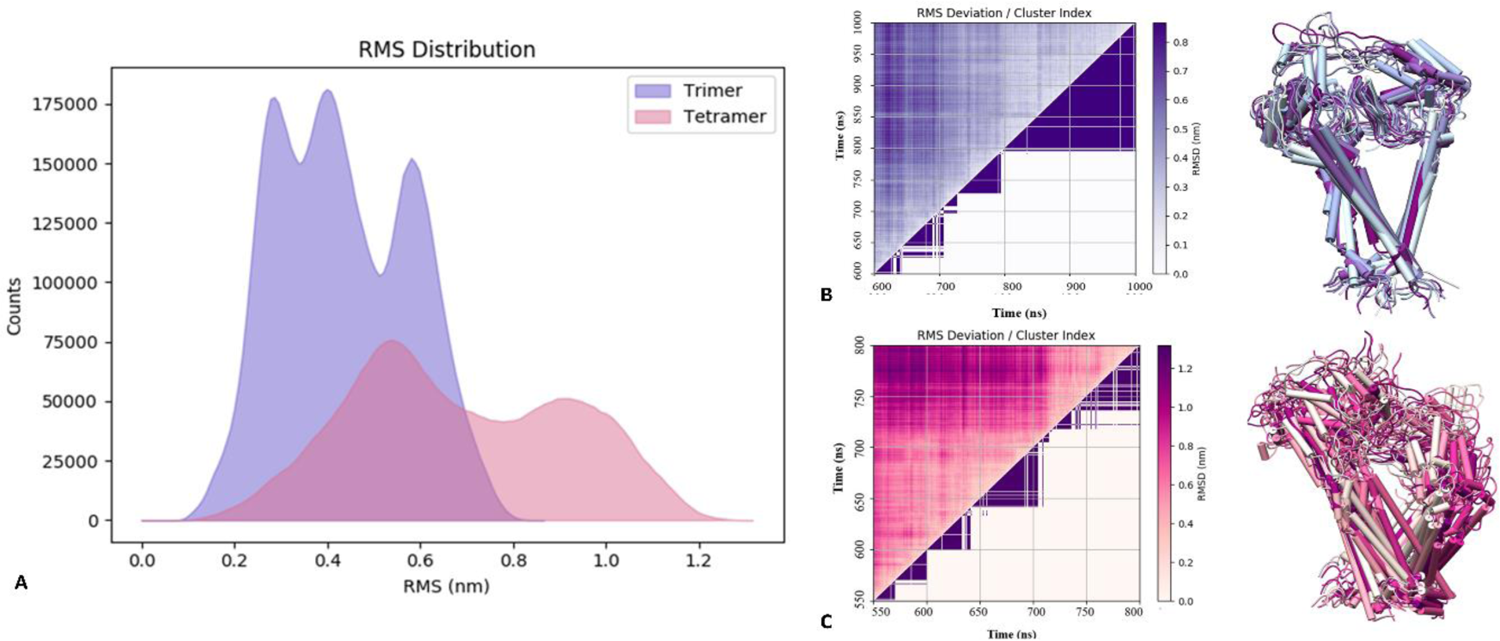
A: RMS Distribution of Trimer and Tetramer Clusters B: Cluster index of Trimer along with cartoon depictions of each cluster representative structure C: Cluster index of Tetramer along with cartoon depictions of each cluster representative structure.

### Stability Check Analysis

It can be deduced from Figure S9; that replicate 8 of the trimer and replicate 2 of the tetramer exhibited quite stable conformations showing high probabilities which were calculated using Kernel Density Estimations (KDE) of root mean square deviations (RMSDs) of the simulation data. In contrast, replicate 6 of the trimer and replicate 8 of the tetramer showed relatively unstable conformations among the evaluated structural ensembles of the respective oligomeric state.

**Figure S9.**
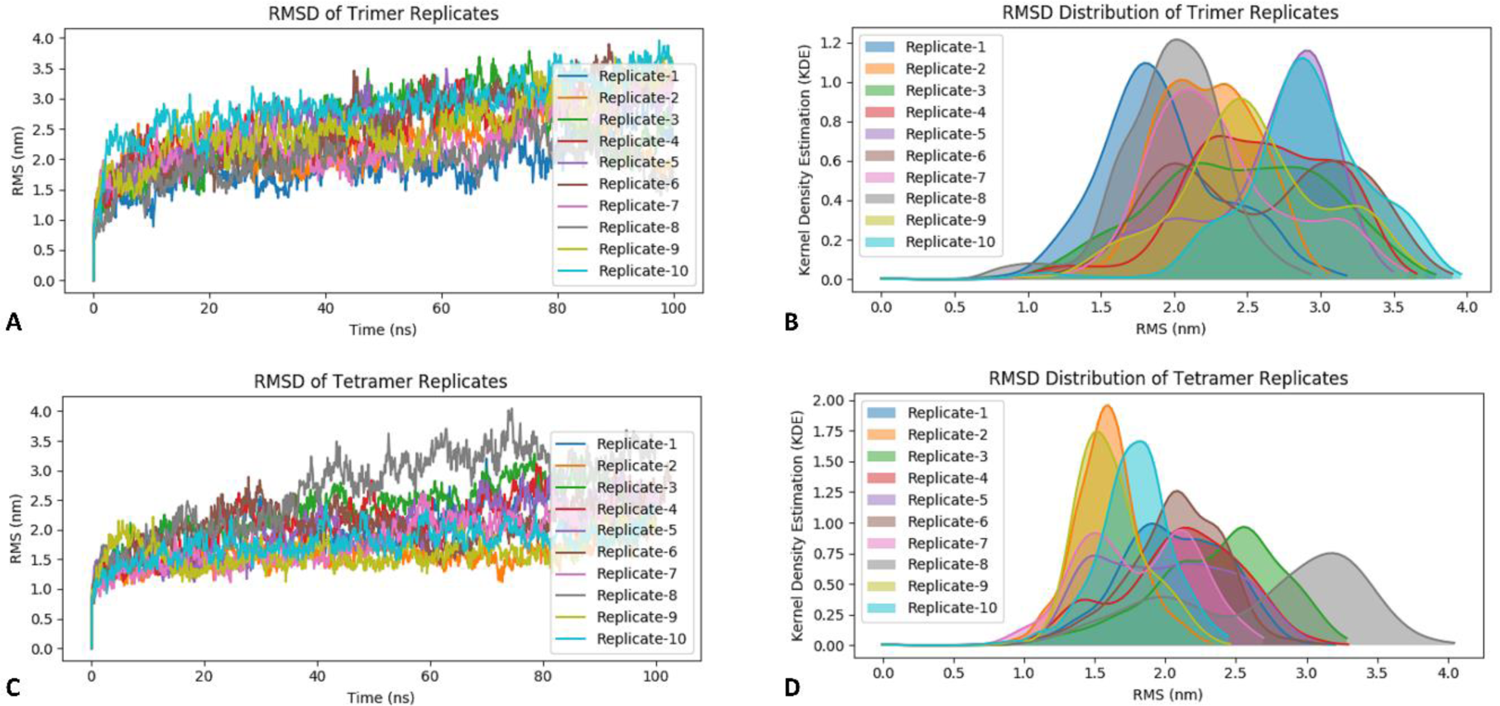
A, C: RMSD of backbone atoms of C-terminal domain of Timer and Tetramer Replicate, respectively along with B, D: their Kernel Density Estimations.

**Figure S10.**
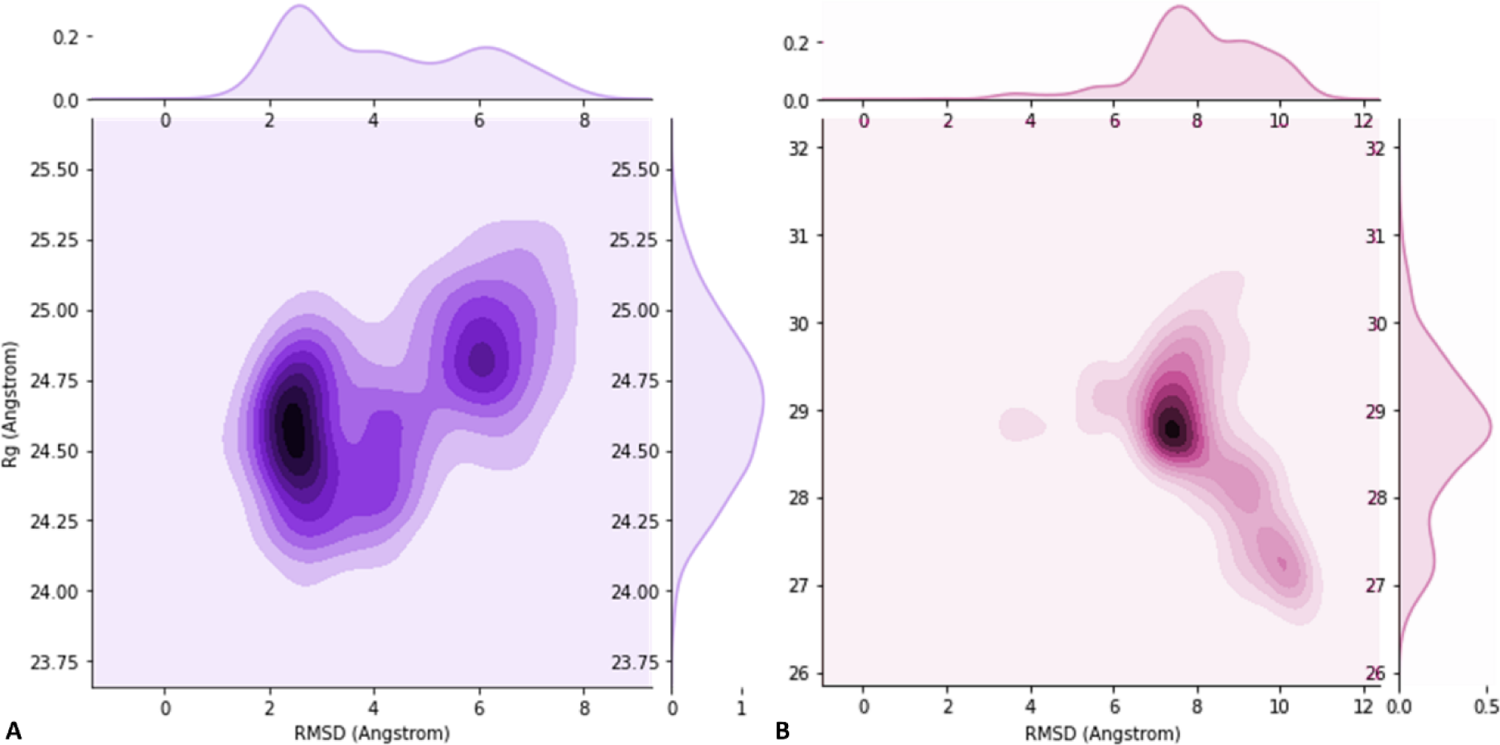
RMSD and R(g) Correlation plot of Trimer **(A)** and Tetramer **(B)**.

The MSD calculations of the oligomeric replicates showed variable dynamical behavior in association with the lipid membrane where replicate 8 of the trimer and replicate 10 of the tetramer maintained the stability along x-y dimensions of the membrane. In contrast, replicate 3 of the trimer and replicate 1 of the tetramer demonstrated exceptional dynamical motions along the membrane planes as evidenced by the plots shown in Figure S11. Moreover, replicate 6 of the trimer also showed some mobility along the membrane plane augmenting the RMSD values, whereas, replicate 8 of the tetramer exhibited a random increase in MSD during the last 40 ns corroborating the RMSD data since an abrupt change was recorded in the stability during the same time frame.

**Figure S11.**
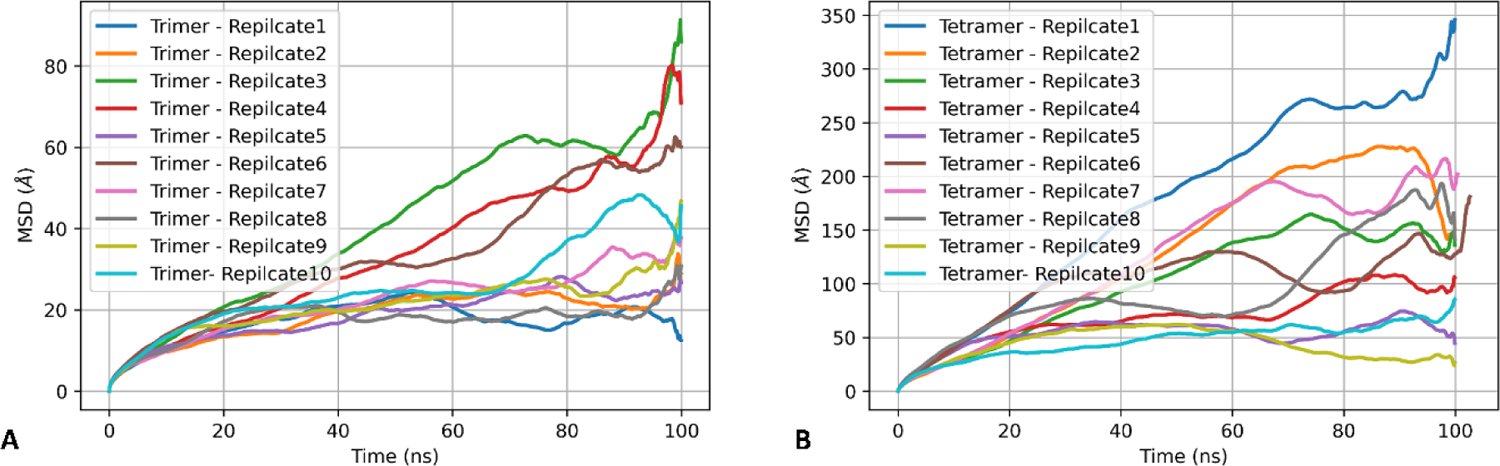
Mean square Displacements of C-alpha atoms of A: Trimer and B: Tetramer along Simulation Times.

**Table S2.** Interaction Occupancies between Basic amino acid residues and lipid molecules.

| Oligomer | Residue Number | Residue Name | Chains | Protein-Lipid Interaction Occupancy (%) |
| --- | --- | --- | --- | --- |
| Trimer | 20 | ARG | A | 46.70 |
|  |  |  | B | 0.00 |
|  |  |  | C | 0.00 |
|  | 29 | LYS | A | 66.26 |
|  |  |  | B | 0.00 |
|  |  |  | C | 92.41 |
|  | 32 | ARG | A | 89.22 |
|  |  |  | B | 0.00 |
|  |  |  | C | 67.46 |
|  | 44 | ARG | A | 0.00 |
|  |  |  | B | 0.00 |
|  |  |  | C | 1.19 |
|  | 53 | LYS | A | 99.20 |
|  |  |  | B | 75.24 |
|  |  |  | C | 48.70 |
|  | 59 | ARG | A | 99.60 |
|  |  |  | B | 99.00 |
|  |  |  | C | 97.40 |
|  | 65 | LYS | A | 67.06 |
|  |  |  | B | 95.20 |
|  |  |  | C | 87.02 |
|  | 73 | ARG | A | 99.80 |
|  |  |  | B | 99.80 |
|  |  |  | C | 99.80 |
|  | 78 | LYS | A | 99.20 |
|  |  |  | B | 99.80 |
|  |  |  | C | 99.60 |
|  | 81 | LYS | A | 95.80 |
|  |  |  | B | 99.60 |
|  |  |  | C | 99.80 |
|  | 85 | ARG | A | 38.12 |
|  |  |  | B | 96.00 |
|  |  |  | C | 94.21 |
| **Tetramer** | 20 | ARG | A | 46.70 |
|  |  |  | B | 0.00 |
|  |  |  | C | 0.00 |
|  | 29 | LYS | A | 66.26 |
|  |  |  | B | 0.00 |
|  |  |  | C | 92.41 |
|  | 32 | ARG | A | 89.22 |
|  |  |  | B | 0.00 |
|  |  |  | C | 67.46 |
|  | 44 | ARG | A | 0.00 |
|  |  |  | B | 0.00 |
|  |  |  | C | 1.19 |
|  | 53 | LYS | A | 99.20 |
|  |  |  | B | 75.24 |
|  |  |  | C | 48.70 |
|  | 59 | ARG | A | 99.60 |
|  |  |  | B | 99.00 |
|  |  |  | C | 97.40 |
|  | 65 | LYS | A | 67.06 |
|  |  |  | B | 95.20 |
|  |  |  | C | 87.02 |
|  | 73 | ARG | A | 99.80 |
|  |  |  | B | 99.80 |
|  |  |  | C | 99.80 |
|  | 78 | LYS | A | 99.20 |
|  |  |  | B | 99.80 |
|  |  |  | C | 99.60 |
|  | 81 | LYS | A | 95.80 |
|  |  |  | B | 99.60 |
|  |  |  | C | 99.80 |
|  | 85 | ARG | A | 38.12 |
|  |  |  | B | 96.00 |
|  |  |  | C | 94.21 |

### Per Residual Contribution

**Figure S12.**
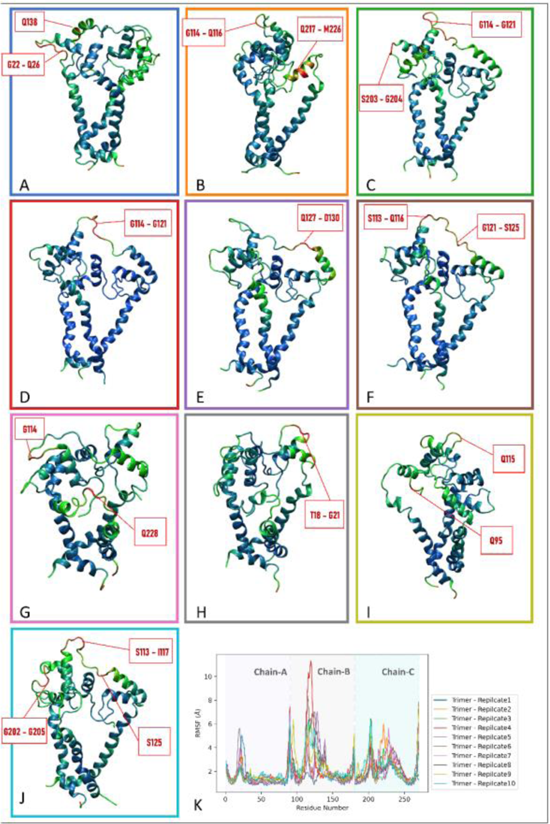
**A-J**: Cartoon representations of trimer from replicate 1 to 10 respectively. Color coding is based on B-factor (Blue to Red), **K**: Root Mean square fluctuation of backbone atoms of trimeric replicates

**Figure S13.**
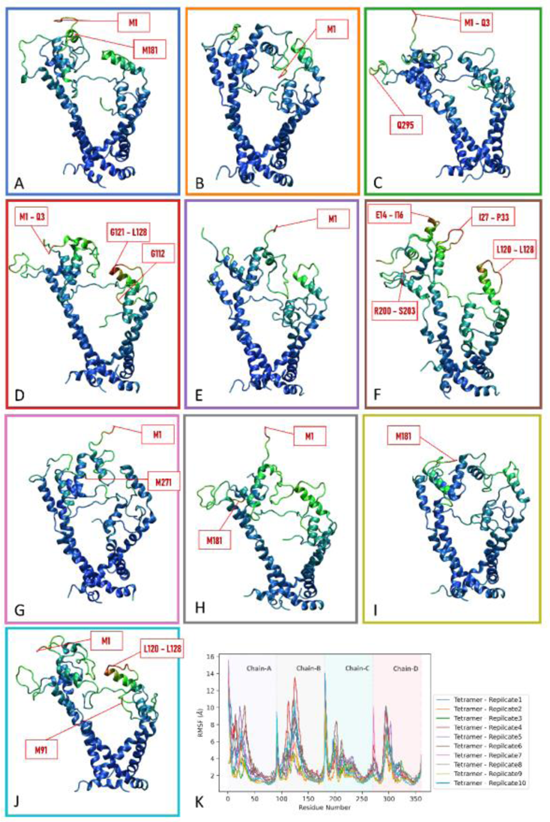
**A-J**: Cartoon representations of tetramer from replicate 1 to 10 respectively. Color coding is based on B-factor (Blue to Red), **K**: Root Mean square fluctuation of backbone atoms of tetrameric replicates.

**Table S3.**
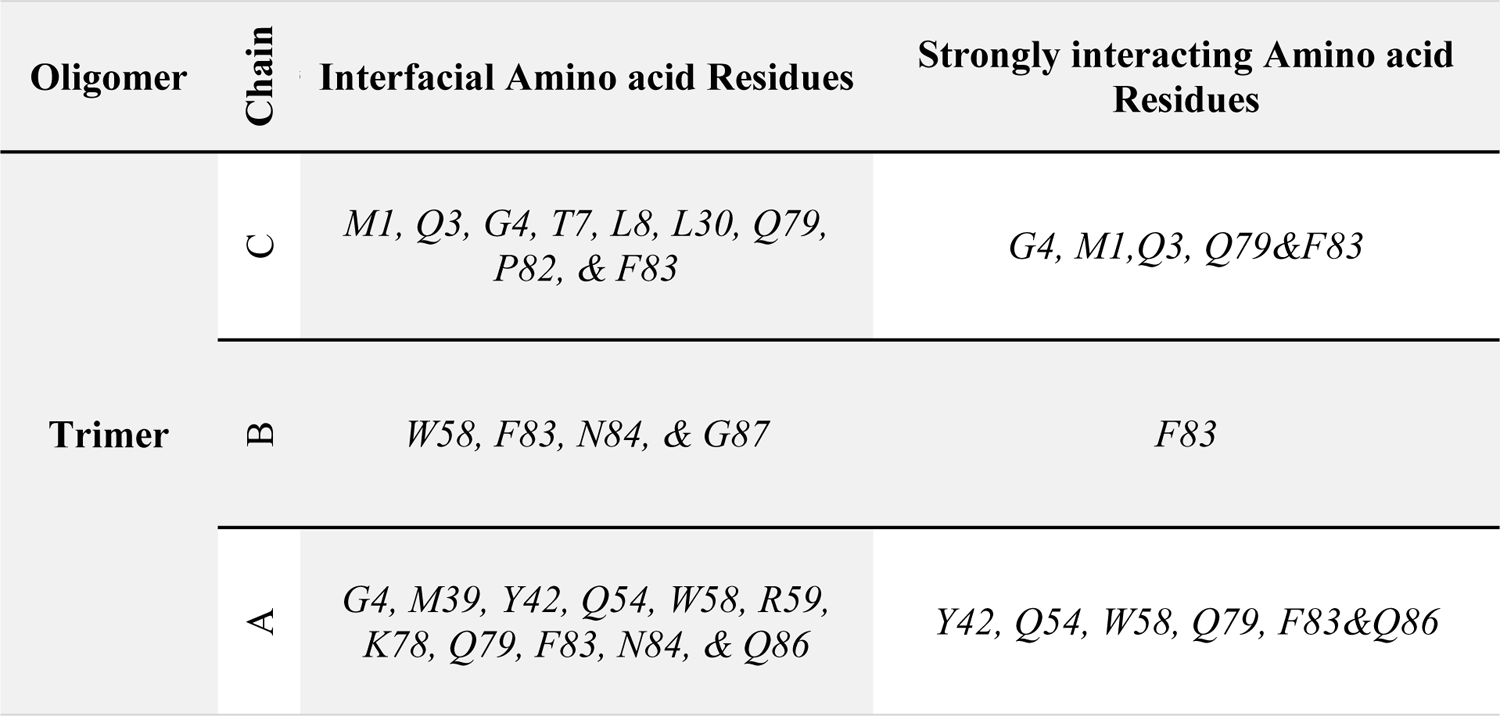
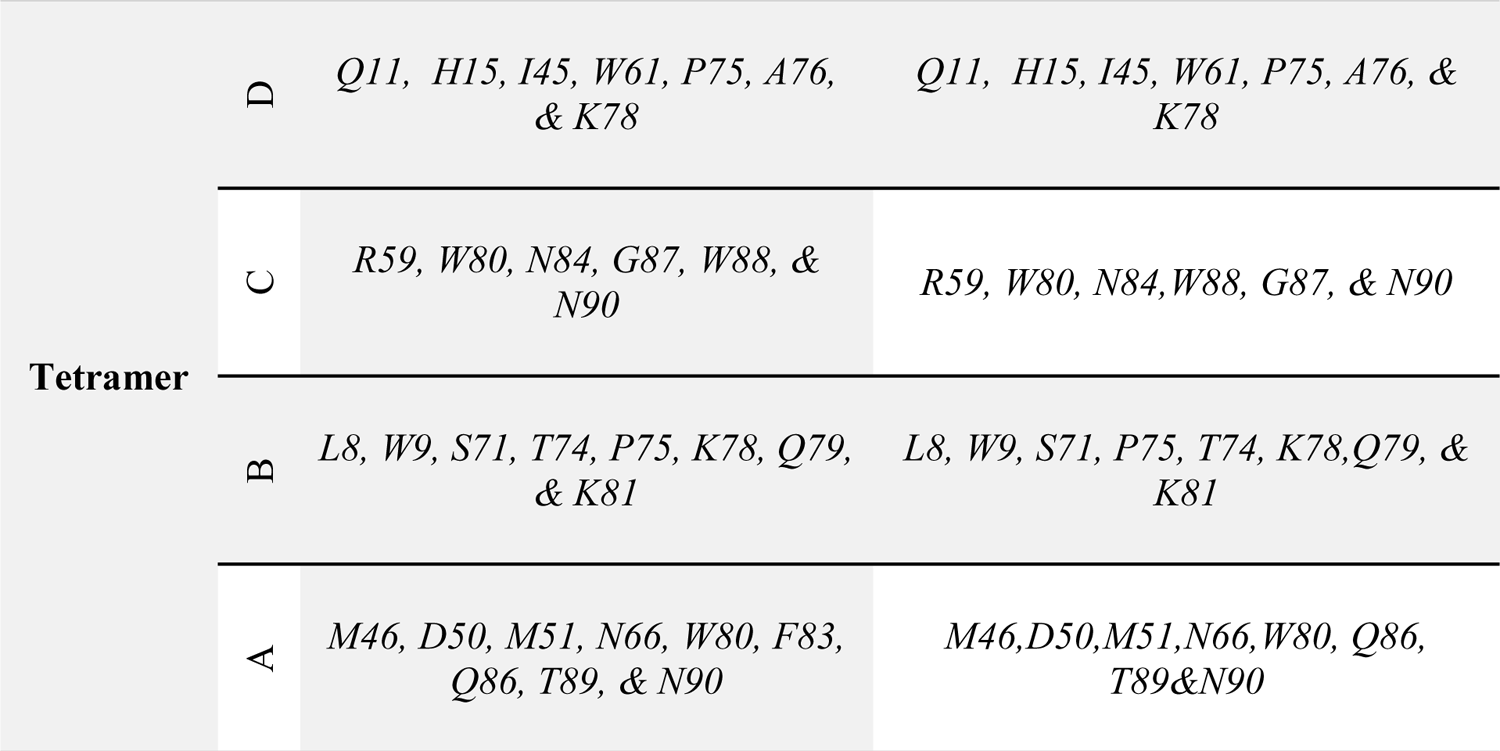
Amino Acid Residues of Trimer and Tetramer involved in interfacial interactions between homo-oligomer chains.

| Oligomer | Chain | Interfacial Amino acid Residues | Strongly interacting Amino acid Residues |
| --- | --- | --- | --- |
| Trimer | C | *M1, Q3, G4, T7, L8, L30, Q79, P82, & F83* | *G4, M1, Q3, Q79 & F83* |
|  | B | *W58, F83, N84, & G87* | *F83* |
|  | A | *G4, M39, Y42, Q54, W58, R59, K78, Q79, F83, N84, & Q86* | *Y42, Q54, W58, Q79, F83 & Q86* |
| **Tetramer** | D | *Q11, H15, I45, W61, P75, A76, & K78* | *Q11, H15, I45, W61, P75, A76, & K78* |
|  | C | *R59, W80, N84, G87, W88, & N90* | *R59, W80, N84, W88, G87, & N90* |
|  | B | *L8, W9, S71, T74, P75, K78, Q79, & K81* | *L8, W9, S71, P75, T74, K78, Q79, & K81* |
|  | A | *M46, D50, M51, N66, W80, F83, Q86, T89, & N90* | *M46, D50, M51, N66, W80, Q86, T89 & N90* |

**Figure S14.**
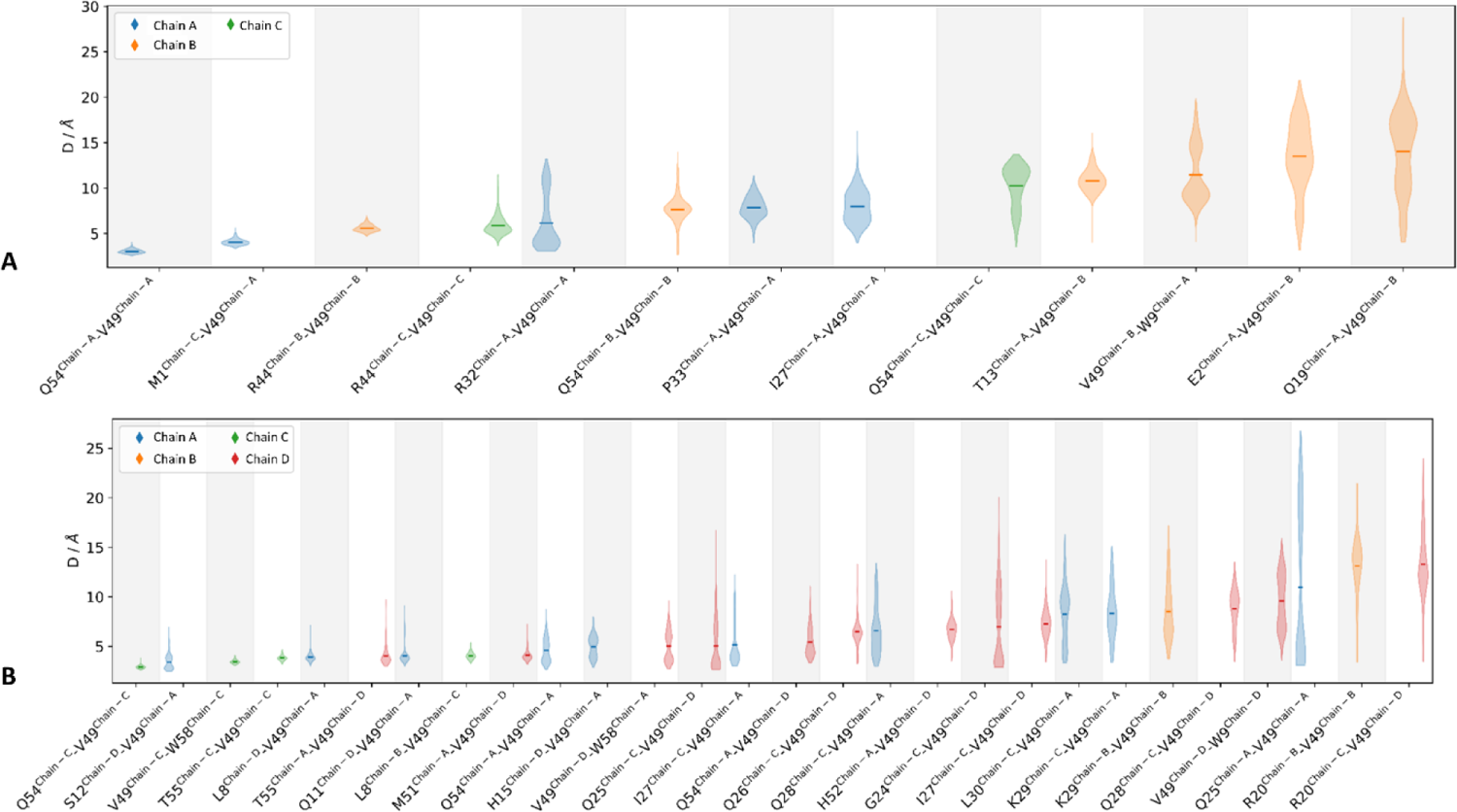
Distance Distributions of V49 neighboring residues of **A**: Trimer and **B**: Tetramer form within a 4 Å cut-off distance

### Characterization of Pore forming Amino Acid Residues

**Figure S15.**
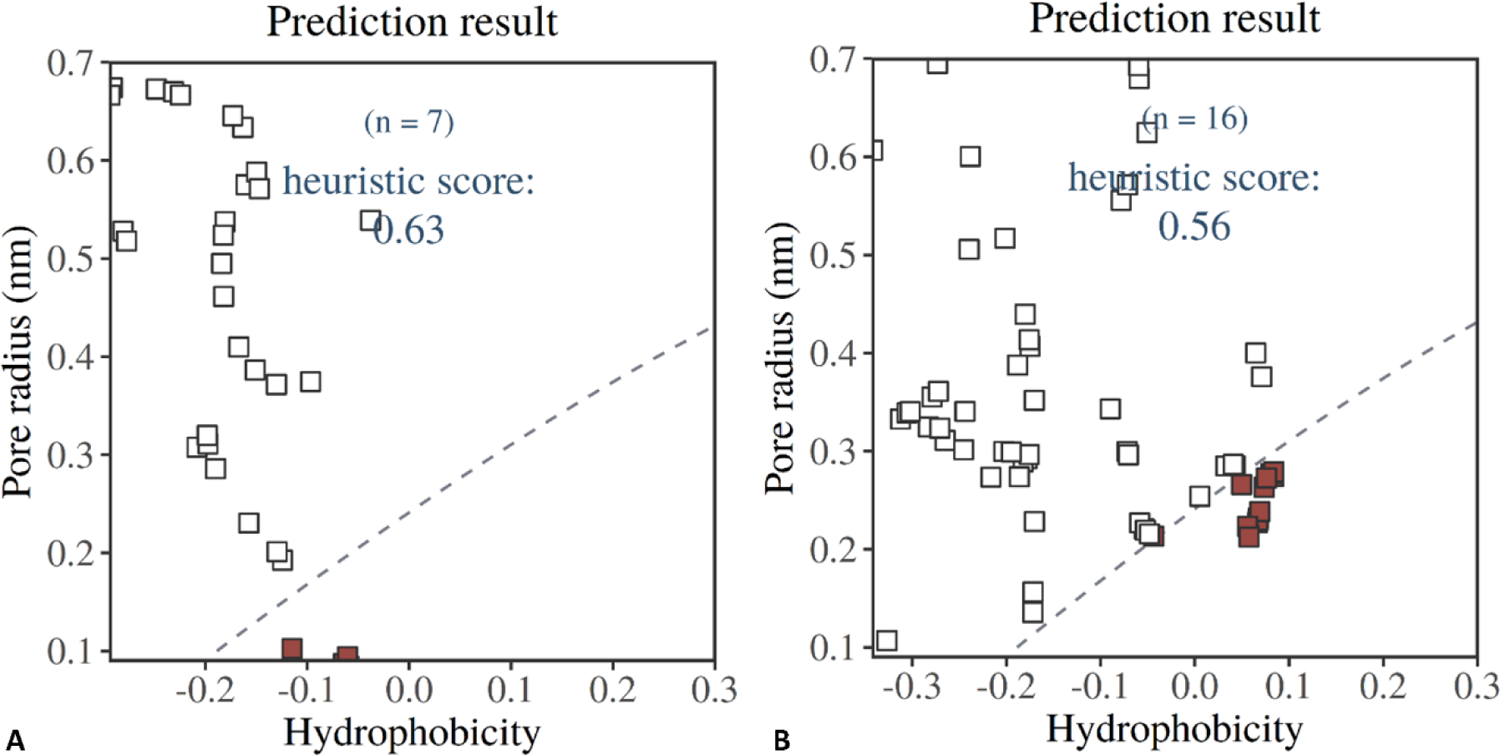
Heuristic Prediction of **A**: Trimer and **B**: Tetramer, where the transmembrane pore radius is plotted on the y-axis against the corresponding local hydrophobicity value of the residue on the x-axis. The dashed line differentiates between the wetted (above the line) and dewetted (below the line) state of the channel as the points (colored red) in the dewetted region predict the closed gates by the summation of the shortest distances between the dashed (1 RT) line and all the red points. If the heuristic score is Σd > 0.55, a channel is said to be in a non-conductive state.

**Table S4:**
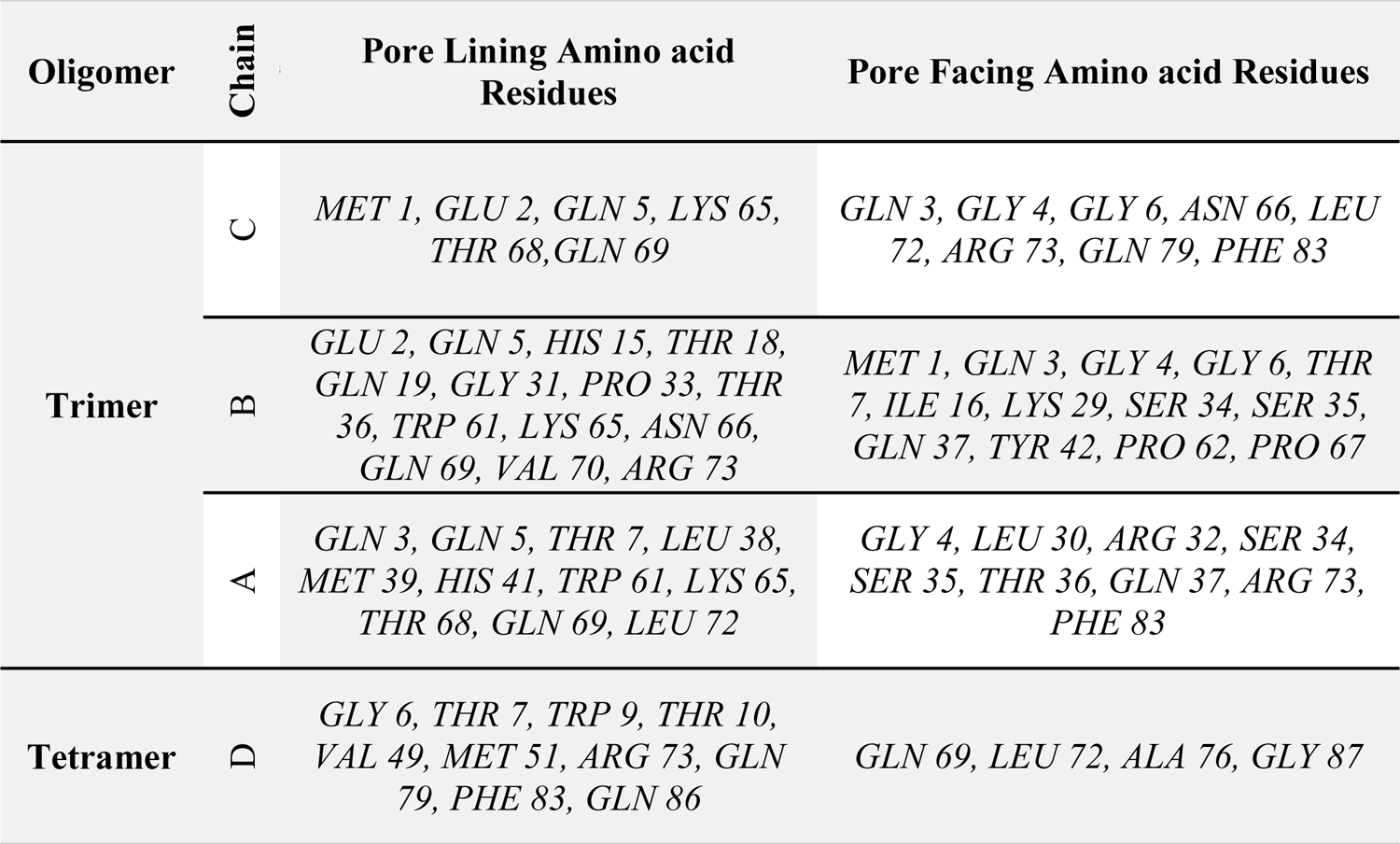
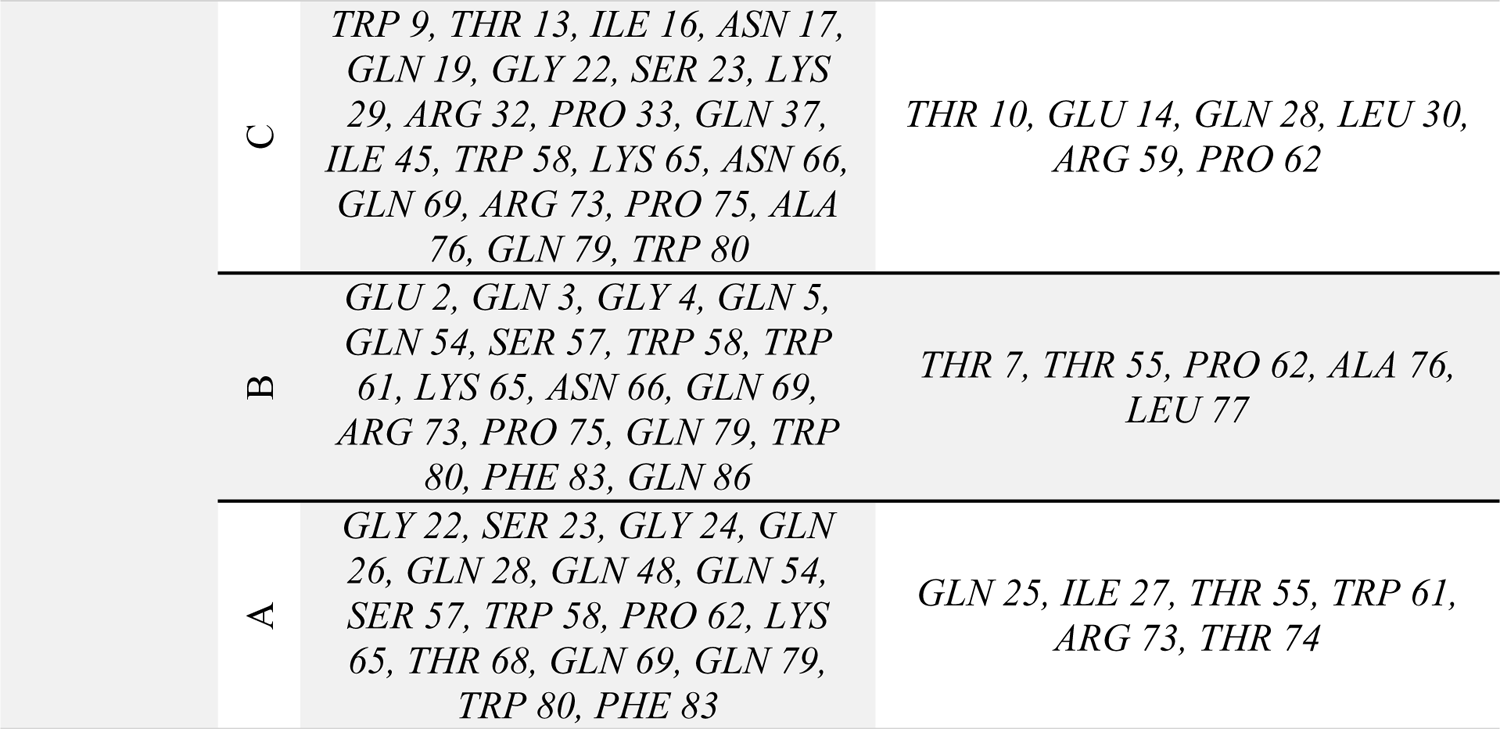
Profiling of Pore lining and Pore facing Amino Acid Residues of Trimer and Tetramer.

| Oligomer | Chain | Pore Lining Amino acid Residues | Pore Facing Amino acid Residues |
| --- | --- | --- | --- |
| Trimer | C | MET 1, GLU 2, GLN 5, LYS 65, THR 68, GLN 69 | GLN 3, GLY 4, GLY 6, ASN 66, LEU 72, ARG 73, GLN 79, PHE 83 |
|  | B | GLU 2, GLN 5, HIS 15, THR 18, GLN 19, GLY 31, PRO 33, THR 36, TRP 61, LYS 65, ASN 66, GLN 69, VAL 70, ARG 73 | MET 1, GLN 3, GLY 4, GLY 6, THR 7, ILE 16, LYS 29, SER 34, SER 35, GLN 37, TYR 42, PRO 62, PRO 67 |
|  | A | GLN 3, GLN 5, THR 7, LEU 38, MET 39, HIS 41, TRP 61, LYS 65, THR 68, GLN 69, LEU 72 | GLY 4, LEU 30, ARG 32, SER 34, SER 35, THR 36, GLN 37, ARG 73, PHE 83 |
| Tetramer | D | GLY 6, THR 7, TRP 9, THR 10, VAL 49, MET 51, ARG 73, GLN 79, PHE 83, GLN 86 | GLN 69, LEU 72, ALA 76, GLY 87 |

|  |  |  |
| --- | --- | --- |
| C | TRP 9, THR 13, ILE 16, ASN 17, GLN 19, GLY 22, SER 23, LYS 29, ARG 32, PRO 33, GLN 37, ILE 45, TRP 58, LYS 65, ASN 66, GLN 69, ARG 73, PRO 75, ALA 76, GLN 79, TRP 80 | THR 10, GLU 14, GLN 28, LEU 30, ARG 59, PRO 62 |
| B | GLU 2, GLN 3, GLY 4, GLN 5, GLN 54, SER 57, TRP 58, TRP 61, LYS 65, ASN 66, GLN 69, ARG 73, PRO 75, GLN 79, TRP 80, PHE 83, GLN 86 | THR 7, THR 55, PRO 62, ALA 76, LEU 77 |
| A | GLY 22, SER 23, GLY 24, GLN 26, GLN 28, GLN 48, GLN 54, SER 57, TRP 58, PRO 62, LYS 65, THR 68, GLN 69, GLN 79, TRP 80, PHE 83 | GLN 25, ILE 27, THR 55, TRP 61, ARG 73, THR 74 |

**Figure S16.**
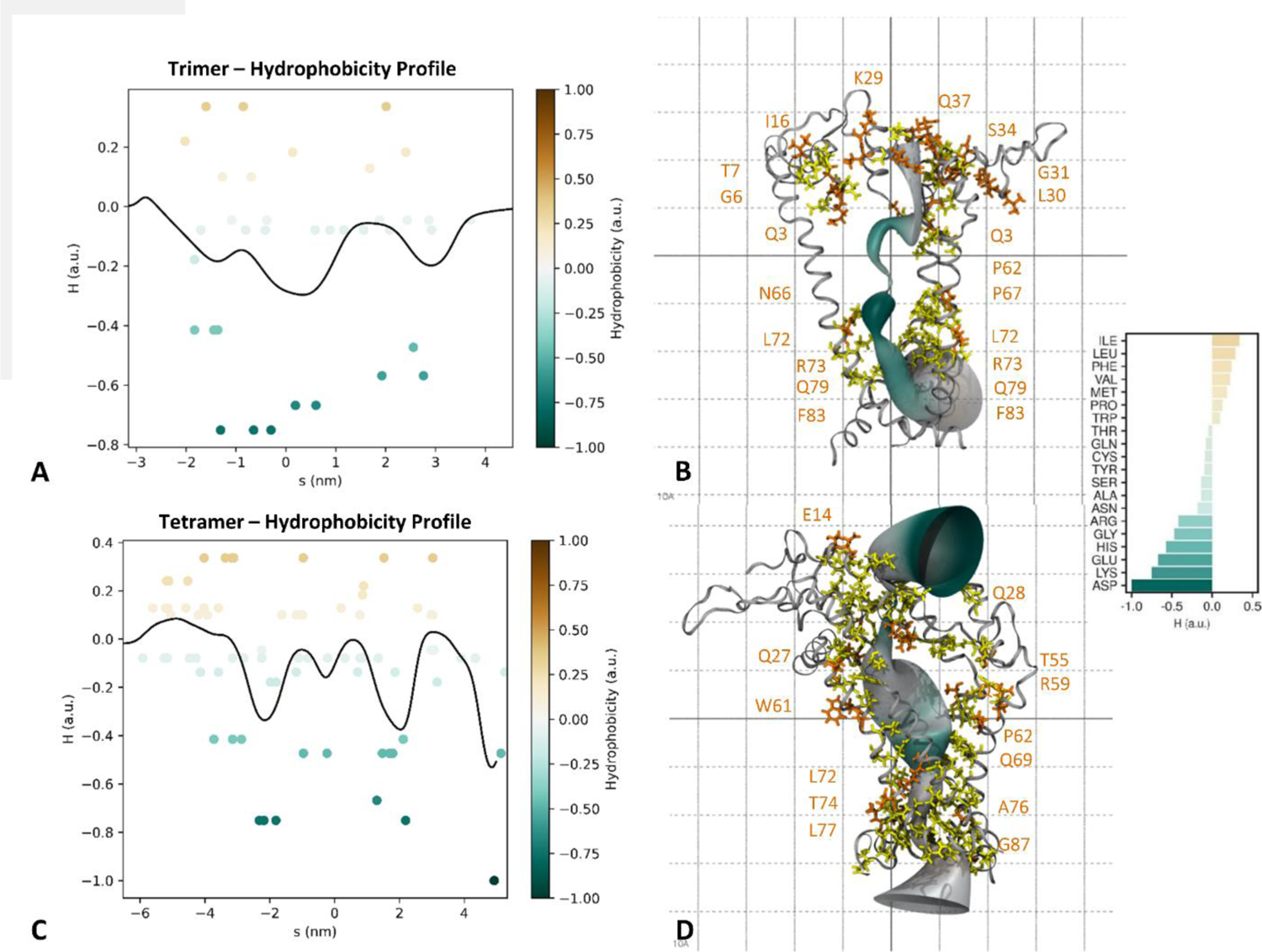
Hydrophobicity profiles of **A**: trimer and **C**: tetramer along with the assignment of hydrophobic surfaces against the pore-lining residues of **B**: trimer and **D**: tetramer (surfaces of brown color contours represent hydrophobic, blue-green are hydrophilic, and white represents neutral amino acids).

**Figure S17.**
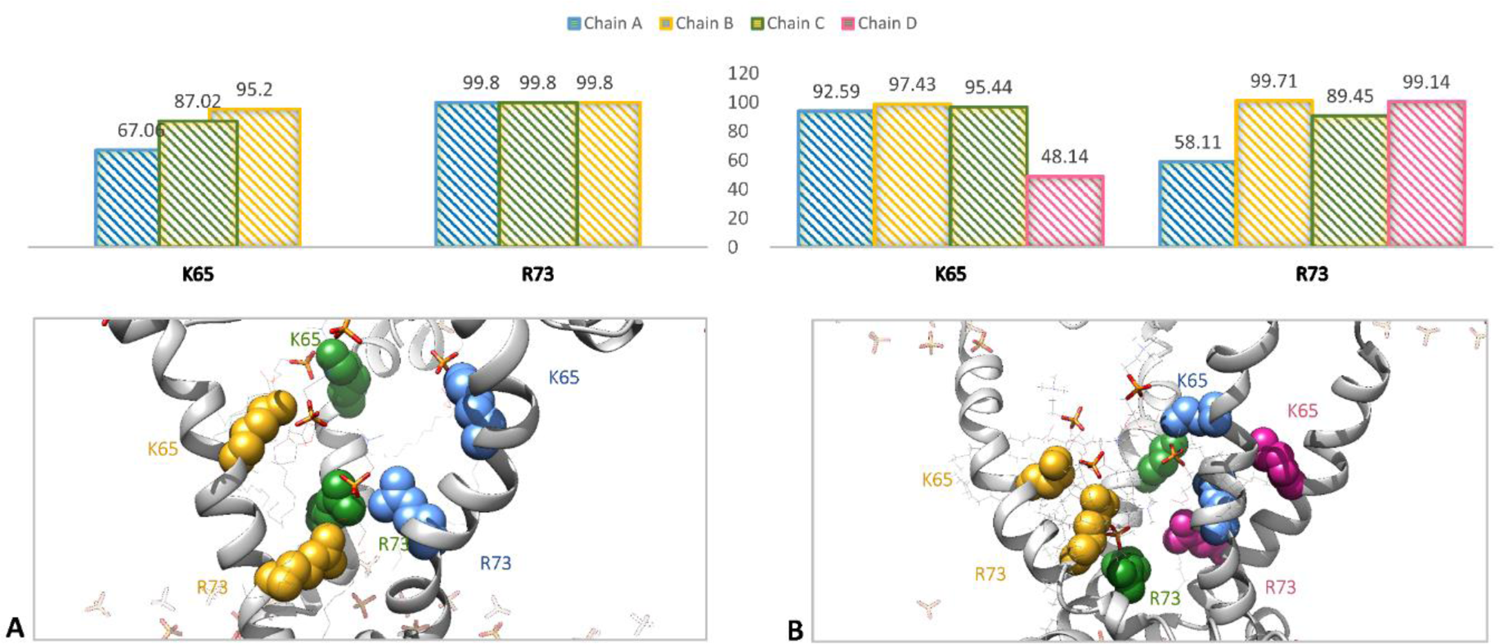
Percentage occupancy of interaction between lipid and cationic amino acid residues inside the pore of the trimeric form (upper left plot) and tetrameric form (upper right plot), along with the cartoon representation of **A**. trimer and **B**. tetramer. The residues of interest (lysine 65 and arginine 73) are depicted as spheres with color coding based on chains A, B, C, and D as blue, yellow, green, and pink, respectively. Lipid phosphate head groups are represented as sticks whereas the hydrophobic side chains are omitted for clarity.

